# Exploration of an enzyme-product mapping approach for plant-derived diterpene synthases

**DOI:** 10.1101/2023.05.26.542545

**Authors:** Yalan Zhao, Yupeng Liang, Xiulin Han, Mengliang Wen

## Abstract

Plant-derived diterpene synthases (PdiTPSs) play a critical role in the formation of structurally and functionally diverse diterpenoids. However, the relationship between PdiTPSs and the specificity or promiscuity of their products remains unclear. To explore this correlation, the sequences of 199 functionally characterized PdiTPSs and their corresponding 3D structures were collected and manually corrected. Using this compiled annotated database, the correlations among PdiTPSs sequences, domains, structures and their corresponding products were comprehensively analyzed. However, utilizing sequence similarity network (SSN), phylogenetic trees, and structural topology features alone was insufficient for effective functional classification of PdiTPSs as these methods could not establish a clear mapping between the enzymes and products. Surprisingly, residues verified to play a function through mutagenesis experiments were located within 8Å of the substrate. Aromatic residues surrounding the substrate exhibited selectivity towards its chemical structure. Specifically, tryptophan (W) was preferentially located around the linear substrate geranylgeranyl pyrophosphate (GGPP), while phenylalanine (F) and tyrosine (Y) were preferentially located around the initial cyclized diterpene intermediate. This analysis revealed the functional space of residues surrounding the substrate of PdiTPSs, most of which have not been experimentally explored. These findings provide guidance for screening specific residues for mutation studies to change the catalytic products of PdiTPSs, allowing us to better understand the correlation between PdiTPSs and their products.

## 1. Introduction

Diterpenoids are a class of widely distributed C20 isoprenoids of natural products, with more than 18,000 members identified in plants (Zeng et al., 2022). They play an important role in plant growth, development (Zerbe and Bohlmann, 2015) and mediate complex plant-environment interactions (Gershenzon and Dudareva, 2007). They also have applications in medicine, flavor, and food industries (Caniard et al., 2012; Jennewein and Croteau, 2001; Philippe et al., 2014; Schalk et al., 2012). All the discovered diterpenoids can be classified according to their core diterpene skeletons by removing all heteroatoms, stereocenters, and reducing unsaturated structures (Bemis and Murcko, 1996; Zeng et al., 2022).

Diterpenoids are highly diversified and complex compounds derived from 5-carbon building blocks isopentenyl pyrophosphate (IPP) and dimethylallyl pyrophosphate (DMAPP). GGPP synthase catalyzes the coupling of IPP and DMAPP in a processive head-to-tail fashion to generate linear hydrocarbon molecules. Then, diterpene synthases (diTPS) and cytochrome P450 monooxygenases (P450s) are responsible for synthesizing a variety of intermediates and modifying skeletons (Banerjee and Hamberger, 2018; Bathe and Tissier, 2019; Chen et al., 2011; Zerbe and Bohlmann, 2015). Particularly, diTPS catalyze remarkably complex cyclization cascades with structural and stereochemical precision and create chemical library of 20-carbon hydrocarbons. Based on the reaction mechanism, diTPS either employ ionization-induced carbocation formation (diTPS I), protonation-induced carbocation formation (diTPS II), or use both mechanisms by bifunctional enzymes (diTPS I/II) (Jia et al., 2018). These diTPS make a significant contribution in synthesizing diverse range of diterpenoid skeletons. However, the limited knowledge on enzyme-substrate recognition and product distribution of diTPS hinder the identification of novel functional diTPS.

Classic multiple sequence alignment methods have been used to identify the functional motifs or sites that affect the product specificity of diTPS. The identified functional motifs include DXDD (Abbas et al., 2017), DDXXD (Rynkiewicz et al., 2001; Starks et al., 1997; Whittington et al., 2002), NSE\DTE (Degenhardt et al., 2009), PIX (Jia et al., 2017) and LHS…PNV (Potter et al., 2014; Potter et al., 2016a; Potter et al., 2016b). By examining the structure-function relationship of *Selaginella moellendorffii* miltiradiene synthase (SmMDS), specific residues around the substrate responsible for product specificity, such as E690, S717, and H721, were identified (Tong et al., 2022). Structural analysis and catalytic mechanism also suggest that the cavity formed by the substrate surrounding residues can selectively choose the substrate (Tao et al., 2022). Product-changing mutational studies and structural analysis provide valuable insights for investigating diTPSs function. However, these researches have only covered a small fraction of the characterized PdiTPSs. To date, no quantitative relationship has been established between sequence and structural features that influence the product specificity of PdiTPSs.

In this study, a manually curated and annotated database has been utilized to investigate the partitioning of PdiTPSs functions using SSN and phylogenetic tree analysis. Then, the correlations were examined among various factors, including overall sequences, subsequences, overall structures, residues around the substrate, and product similarity. Lastly, we calculated the residue preferences surrounding the substrate and analyzed their spatial conservation to determine the range of residues that significantly affect substrate type and product outcome. The results of the comprehensive analysis provide valuable insights into exploring product-specific residues in PdiTPSs and mapping patterns between PdiTPSs and their functions.

## 2. Results and discussion

### 2.1 Overview of functional annotations of PdiTPSs

A manually curated database of 199 functional characterized PdiTPSs has been presented, including 27 bifunctional enzymes, 91 class I enzymes, and 82 class II enzymes (Supplementary Table S1). These PdiTPSs were derived from 69 plant species belonging to 26 families and 52 genera, producing 16 diterpene intermediates and 63 diterpene precursors. Of these products, only a small fraction were found to be associated with multiple PdiTPSs, while the majority of products were primarily catalyzed by a single PdiTPSs, which affected the product-specific analysis.

To solve this problem, the existing terpenoid skeleton classification system (Hu et al., 2021) was employed to group these products into 16 different types. Products from PdiTPS I and PdiTPS I/II were classified into 15 skeleton types, while those from PdiTPS II were classified into only one single type. The classification scheme allowed us to group multiple products from a single enzyme into the same category, such as SsSS synthase from *Salvia sclarea* (Caniard et al., 2012), which could catalyze the dephosphorylation and minor rearrangement of 9 diterpene intermediates to produce 11 diterpene precursors, all of which fall into the same Labdane scaffold (SK1). However, this classification system is not always effective, for example, TrTPS13 from *Tripterygium regelii* produces five products belonging to SK4, SK5, and SK9 scaffolds (ntkrn, sndarpardn, ipsfdn, spsfdn, sdmon), and PdiTPS from *Grindelia hirsutula* produces three products (abedn, epmnlo, mnlo) belonging to SK1 and SK3 scaffolds, respectively.

### 2.2 The sequence similarity network generates PdiTPSs clusters

The results of applying a skeleton classification system for functional mapping in SSN analysis was examined. This all-pairs local sequence-based comparison method could rapidly generate a network of nodes and edges using any expectation value (E) as a threshold. By appending annotation information, the sequence-function relationship profile of the enzyme could be quickly viewed.

This analysis results suggested that C-terminal, N-terminal, NC terminal subsequences and overall sequences could be used to classify the PdiTPS I and PdiTPS II. In general, the N-terminal (Fig. 1A), C-terminal (Fig. 1B) and NC-terminal (Fig. 1C) networks generated by SSN resulted in mainly multiple backbone clustering. In particular, SK1, SK3, and SK4 skeletons were often clustered together, despite the clear differences in their product structures. In contrast, the product skeleton clusters obtained from multiple sequence comparison of N-terminal subsequences were more refined, with fewer outliers. Multiple sequence comparisons also revealed that PdiTPSs from different species were clustered together, which might limit the grouping of PdiTPSs by product type. Additionally, it could be observed that products belonging to the SK1, SK3, SK4, and SK5 skeletons are predominantly found in early diverging plant lineages such as ferns and mosses (Fig. 1). These skeletons, including labdane (SK1), abietane (SK3), pimarane (SK4), and kaurane (SK5), are the most abundant and widely distributed (Johnson et al., 2019). SSN analysis yielded additional insights into the relationship between product backbone and enzyme sequence (Fig. 1C). Specifically, our results suggest that the SK1 and SK5 backbones serve as crucial linkage points for other backbone groups. It is important to note that while the SK1 and SK5 skeletons may serve as connection points for other skeleton clusters, this does not necessarily imply that they are the fundamental skeletons driving the evolution and diversification of diterpene synthase products. Further experimental research is needed to confirm this.

**Fig. 1.**
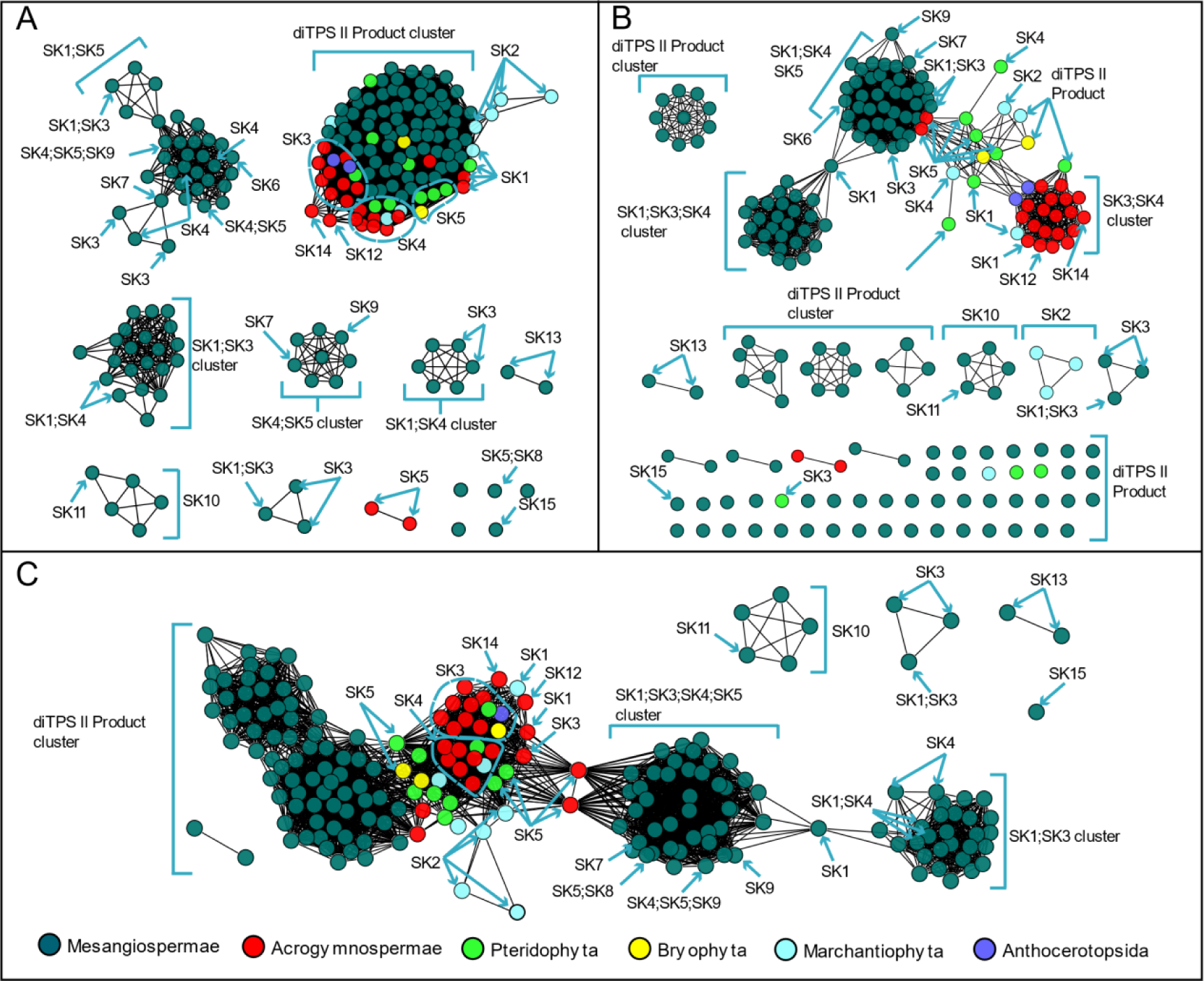
Similarity networks of PdiTPSs sequences. A) Clusters of PdiTPSs product skeletons defined by N-terminal subsequence similarity (E-value threshold 10^-70^); B) Clusters of PdiTPSs product skeletons defined by C-terminal subsequence similarity (E-value threshold 10^-70^); C) Clusters of PdiTPSs product skeletons defined by NC-terminal subsequence similarity (E-value threshold 10^-120^).

### 2.3 PdiTPSs products evolve to polycyclic structures

Multiple sequence alignment and evolutionary information in phylogenetic analysis can be used for comparing protein homology, providing insight into protein sequences, domains and motifs, specific conserved sites evolution, and functional variation. The phylogenetic tree of PdiTPSs was constructed to detect evolutionary relationships and identify lineages with similar features. By examining the changes in product skeleton, it could be able to identify potential correlations between product and PdiTPSs mapping. Terpene synthases commonly contain two conserved structural domains, the N-terminal and C-terminal domains. Therefore, it was also constructed for the phylogenetic trees for N-terminal, and C-terminal subsequences.

Unfortunately, the phylogenetic tree did not provide a clear division of PdiTPSs based on their functions. However, it has been found that SK1, SK2, SK5, SK4, and PdiTPS II-products were frequently present in the early diverging PdiTPSs products. This rule can be indicated by the phylogenetic trees of overall sequences (Fig. 2), N-terminal (Supplementary Fig. S1A) and C-terminal (Supplementary Fig. S1B) domains. On the other hand, SK6, SK7, SK 8, and SK9 were found in the late-emerging PdiTPSs products. It could also be observed for the trend of PdiTPSs product functions evolving towards multi-ring skeletons from the major branches of the trees. Moreover, the PdiTPSs that produced the SK10, SK11, SK12, SK13, and SK14 skeletons showed shorter evolutionary distances from the ancestral PdiTPSs. This provides valuable insights into how the function and evolution of PdiTPSs may contribute to species-specific adaptations to unique ecological niches. However, additional investigations are necessary to further explore the distribution patterns of diterpenoid compound types and their relationship with the evolutionary status of plants. Similar research has been carried out to investigate the distribution of terpenoid compounds and their biosynthetic pathways in various species of *Isodon* plants (Li et al., 2013).

**Fig. 2.**
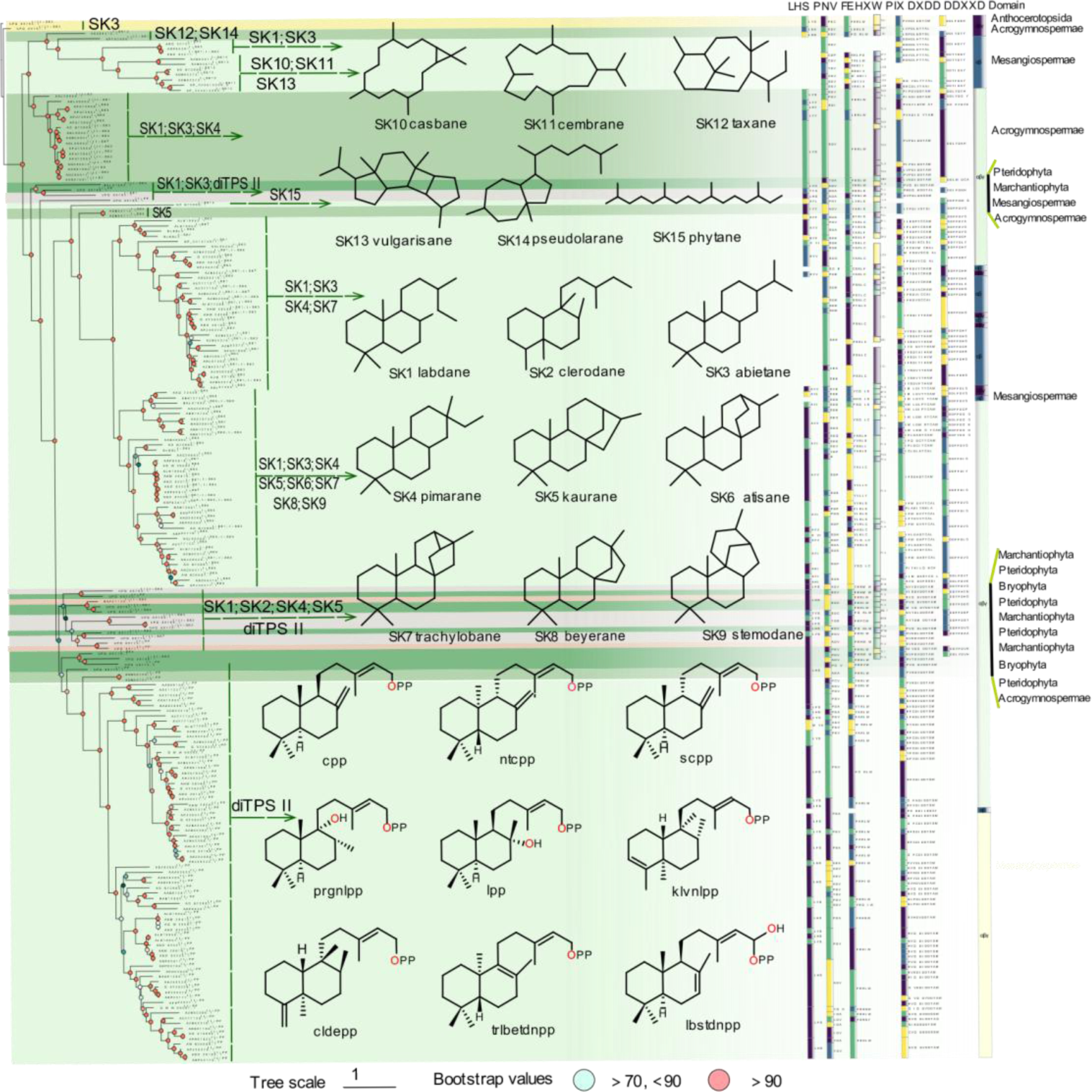
The relationship between the overall sequence phylogeny of PdiTPSs, their product scaffolds, function-related motifs, and enzyme source classification. The phylogenetic tree labels each enzyme with its accession ID, product class, and scaffold classification. It also displays the skeleton structures of SK1-SK15, major product structures of diTPS II, function-verified motifs in some diterpene synthases, and the domain composition of each diterpene synthase.

Our analysis of the domain composition of PdiTPSs showed that the *γβα* triple-domain structure and *βα* bi-domain structure alternately appeared in the phylogenetic tree, indicating a phenomenon of continual loss and acquisition of structural domain subsequences during the evolution of terpene synthases. In addition, the bi-domain *βα* structure was only found in the angiosperms (Fig. 2), supporting the origin of the *βα* bi-domain structure, which was generated by the loss of the *γ* domain in ancestral terpene synthases that had the *γβα* structure (Christianson, 2017; Wang et al., 2023).

The LHS and PNV motifs (Fig. 2) were CPS-specific motifs (Potter et al., 2014), as confirmed by our computational analysis. These two motifs were conserved in PdiTPS II, but had undergone mutations in PdiTPS I. The histidine (H) residue in the FEHXW motif exerted cooperative GGPP/Mg^2+^ inhibition on CPS (Jia et al., 2022), but histidine was not always conserved in the FEHXW motif of PdiTPS II. Although the function of aromatic amino acids in this motif remained unclear, it had been observed in PdiTPS I that these residues were no longer predominantly composed of aromatic amino acids in this motif, but rather of aliphatic and uncharged amino acids. The PIX motif (Fig. 2) displayed was related to *ent*-kaurene synthesis (Jia et al., 2017), and was lost in the PdiTPSs of angiosperms that produced primarily SK1 and SK3, as well as in the that of polycyclic skeleton SK10, SK11, and SK13. This motif was present in the PdiTPSs of mosses that produce SK1 and SK3, but had undergone mutations. This shows that product-specific motifs can exist in PdiTPSs, which have evolved through deletions and mutations that have resulted in enzyme sequences different from their ancestors and acquisition of new functions. Therefore, motifs that are different from the ancestral enzyme and absent in other PdiTPSs may help uncover product-specific motifs.

### 2.4 Identify conserved and diverse subsequences in PdiTPSs via sequence similarity analysis

The sequence similarity features of PdiTPSs were examined, where the larger the upper quartile and lower quartile in the box plot, the higher the sequence similarity. Statistical analysis (Fig. 3A) showed that the C-terminal was more conserved in PdiTPS I, while the N-terminal was more conserved in PdiTPS II. The main differences in sequence similarity between PdiTPS I and PdiTPS II were located in the C-terminal (Fig. 3A), suggesting that the use of C-terminal subsequences might facilitate the divergence. The similarity distribution of PdiTPS I and PdiTPS I/II sequences was lower in the NC-terminus and overall regions than that of in the C- and N-terminal subsequences, while the similarity distribution of PdiTPS II and PdiTPS I/II sequences was lower in the C-terminus than that of in the N-, NC-, and overall regions (Fig. 3A). Therefore, when distinguishing between PdiTPS I and PdiTPS I/II, it was necessary to consider the differences between the NC-terminus and overall sequences, while differences in the C-terminal subsequence might be helpful for distinguishing PdiTPS II from PdiTPS I/II. Additionally, it should be noted that PdiTPS I/II was relatively conserved in the N-, C-, NC-, and overall sequences, with sequence similarities mostly above 50%, especially in the N-terminal subsequence, which exhibited the highest fourth quartile of sequence similarity values. Thus, the N-terminal subsequence might be useful for clustering PdiTPS I/II.

**Fig. 3.**
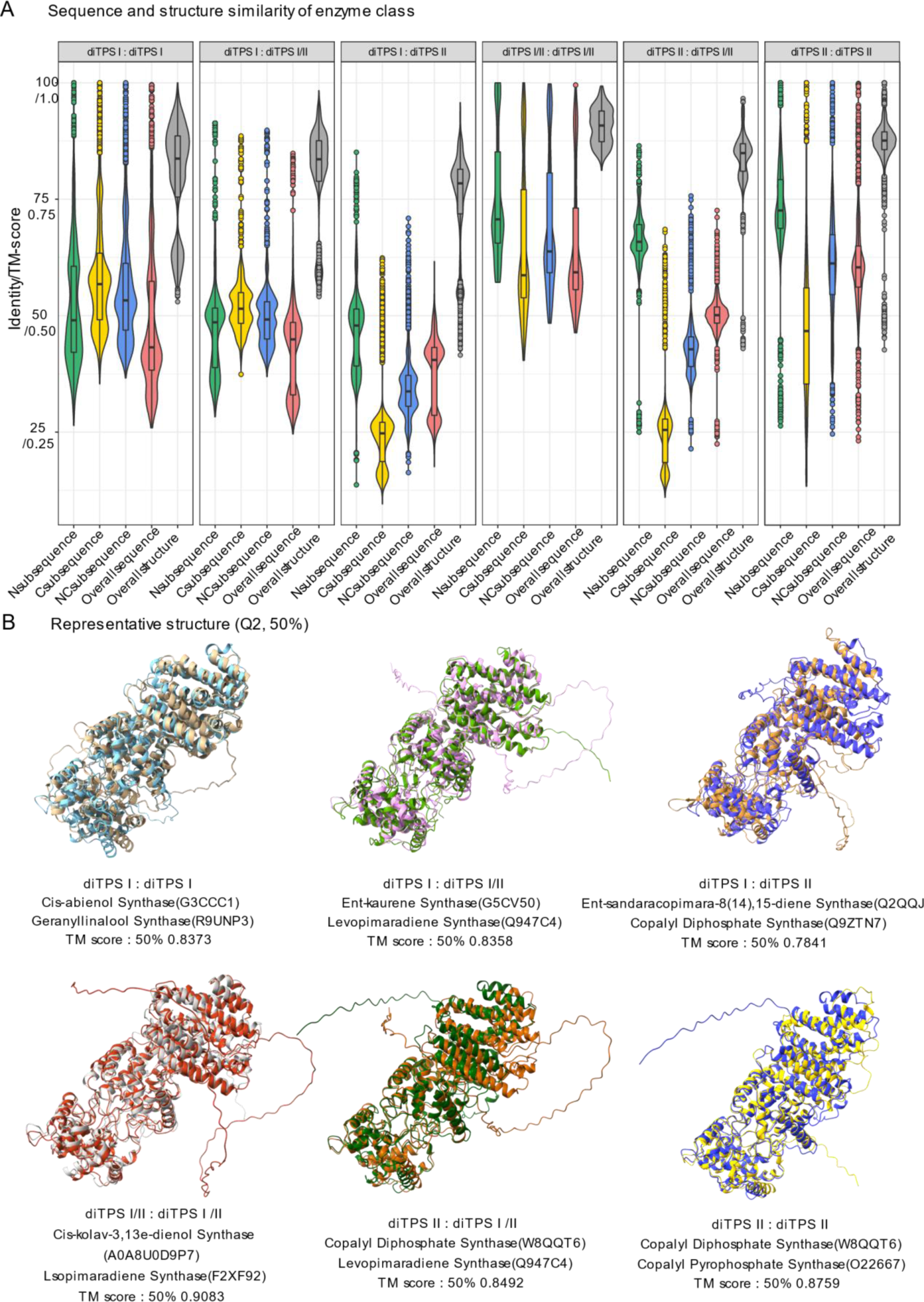
Distribution of sequence and structural similarity data. A) Distribution of sequence and structural similarity data among N-terminal, C-terminal, NC-terminal domains, overall sequences, and overall structures of diTPS I, II, and I/II. B) Superimposition of representative structures at the Q2 position based on the TM-score.

After obtaining the basic conservation features of the PdiTPSs sequences, it had been comparatively analyzed for the sequence similarity in accepting the same or different substrates and producing the same or different products. Theoretically, the sequence similarity that recognize the same substrates or produce the same products should be higher than those that recognize different substrates or produce different products. The results showed that the N-terminal subsequence had the highest upper and lower quartiles of sequence similarity in identifying the same substrate and producing the same product, while the C-terminal subsequence had the lowest lower quartile of sequence similarity in identifying different substrates and producing different products. Additionally, the C-terminal subsequence had the fewest 100% sequence similarity values in identifying different substrates and producing different products (Figs. 4A and 4B).

**Fig. 4.**
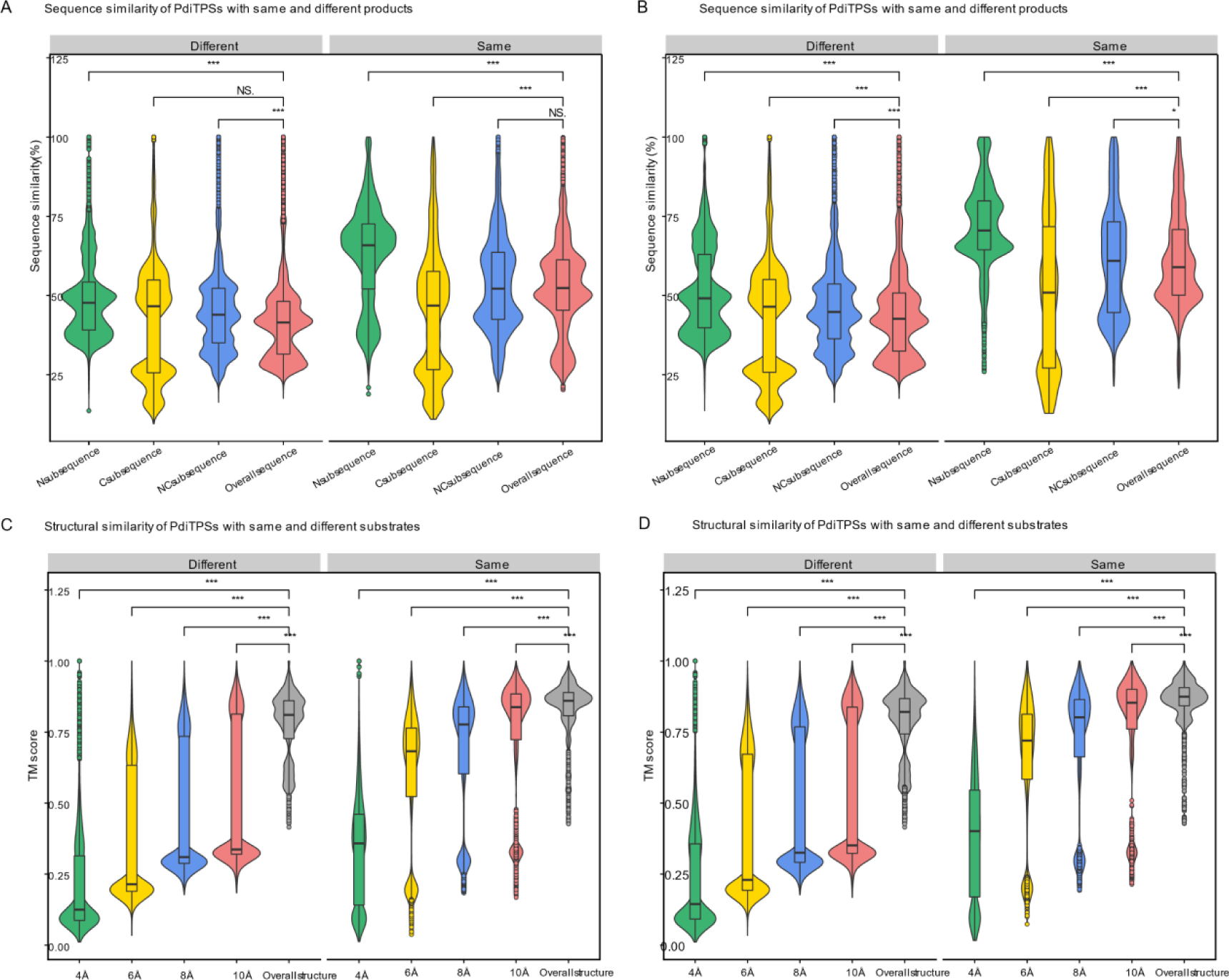
Distribution of similarity data between N-terminal, C-terminal, NC-terminal subsequences, overall sequences, and overall structures of PdiTPSs for same and different substrates and products. A) Comparison of sequence similarity data between N-terminal, C-terminal, and NC-terminal subsequences and overall sequences for the same and different substrates. B) Comparison of sequence similarity data between N-terminal, C-terminal, and NC-terminal subsequences and overall sequences for the same and different products. C) Comparison of sequence similarity data between the topology structures formed by residues within 4 Å, 6 Å, 8 Å, and 10 Å of the substrate with the overall structures for the same and different substrates. D) Comparison of sequence similarity data between the topology structures formed by residues within 4 Å, 6 Å, 8 Å, and 10 Å of the substrate with the overall structures for the same and different products. Asterisks indicate statistical significance (*: p < 0.05, **: p < 0.01, ***: p < 0.001), and “NS” indicates no statistical difference.

Similar results have been reported in the study of sesquiterpene synthases, where the phylogenetic tree constructed using the C-terminal subsequence can group enzymes based on their product types, and the addition of the N-terminal subsequence has little effect on the structure of tree (Durairaj et al., 2019). This is also reflected in our SSN analysis, where the C-terminal subsequence generated more different clusters and distant outliers for different types of products than the N-terminal subsequence at the same threshold (Figs. 1A and 1B).

### 2.5 PdiTPSs have a conservative structure, but the surrounding residue topology is flexible

Through sequence analysis, some clues had been obtained to predict the substrates and products of PdiTPSs, but more evidence still was needed to help us predict the products. Therefore, it would be needed to further examine the structure of PdiTPSs and their residues around the substrates. AlphaFold2 was applied to help rebuild the structures of all collected PdiTPSs. Most amino acid residues had a high credible score for pLDDT, indicating that the predicted structures were highly accurate and close to X-ray resolutions.

It had been calculated for the distribution of topological similarity of the overall structure (Fig. 3A) and the TM scores for topological similarity were mostly greater than 0.75. Furthermore, the median TM score of the overall structure when recognizing the same substrate and different substrates (Fig. 4C), as well as producing the same and different products (Fig. 4D), were also greater than 0.75 and closer to the upper quartile (Q3). These results suggested that PdiTPSs that perform different functions shared a similar TPS fold. Here, only the superimposed results of the representative structures of different types of PdiTPSs at the median (Q2) of TM score were shown (Fig. 3B). Supplementary Fig. S2 showed the structural superposition results of the representative structures of different types of structures representing PdiTPSs in Fig. 4 with 2 extreme values and 3 quartiles of TM score. In addition, as the selected residue range around the substrate increased, the TM score also increased, indicating that the difference of topological structure formed by residues closer to the substrate was greater, while the conservation of the topological structure formed by residues further away from the substrate increased, but the increasing trend became flat (Figs. 4C and 4D).

Both the structure formed by substrate-surrounding residues and the overall structure were significantly higher distributed in the TM score of the same substrate or same product than those of the different substrates and different products (Figs. 4C and 4D). This trend was more significant than the overall difference trend contributed by sequence similarity (Figs. 4A and 4B). Similar to the evaluation of sequence similarity, it was expected for the TM score between substrate-surrounding residues and overall structures that recognized the same substrate or produced the same product to be TM score > 0.5, and TM score of those that recognized different substrates and produced different products to be TM score < 0.5 in the analysis of structures. Therefore, the topological structures formed by residues within 6Å of the substrate appeared to be the best choice for determining substrate and product similarity.

### 2.6 N-terminal subsequence strongly correlates with overall sequence similarity

The above results have provided some insights for the determination of product types at both the sequence and structure levels. Hence, it would be interested to further explore methods for quantitative assessment the relationship between PdiTPSs sequences, structures and products. The correlation was evaluated using Pearson’s correlation coefficient (PCC) (Riziotis et al., 2022), with only the final coefficient shown here. The statistical results showed that the similarity between the C-terminal subsequence and the overall sequence (PCC = 0.46, *p* < 0.001) was significantly weaker than that between the N-terminal subsequence (PCC = 0.91, *p* < 0.001). And there was almost no correlation between the C-terminal and N-terminal subsequences (PCC = 0.26, *p* < 0.001).

The weak correlation in sequence similarity between these two domains supports their independent origins. However, an intriguing observation is that the sequence similarity between the N-terminal subsequence and the full-length sequence is highly correlated, while the C-terminal subsequence is not correlated with either the N-terminal or full-length sequences. Nevertheless, the similar phylogenetic tree structures constructed from the C-terminal subsequence, the N-terminal subsequence, and the full-length sequence suggest that although the C-terminal and N-terminal subsequences originated independently, they have co-evolved. During this process, the N-terminal subsequence underwent strict purification selection and evolved at a slow rate, as indicated by the lack of catalytic activity of the *β*-domain in Class I diterpene synthases, which has been conserved during evolution (Faylo et al., 2021). In contrast, the C-terminal subsequence underwent functional selection and evolved at a faster rate to acquire new functions to adapt to the environment. Clues to this can be found in our statistics of the sequence divergence of the C-terminal subsequences than the N-terminal subsequences.

### 2.7 Substrate-surrounding residue topology in PdiTPSs is independent of the overall structure

Obviously, the analysis based on sequence traits provide limited insights into substrate recognition and functional diversification of PdiTPSs. Protein structures provide a higher resolution platform for understanding function, but acquiring protein structures is expensive and difficult. Therefore, the crystal structures of diterpene synthases are also limited, and the emergence of AlphaFold2 and its high accuracy is exciting, as it has been applied to understand the mechanisms of enzyme (Zhai et al., 2022). AlphaFold2 has been applied to obtain structural data for the PdiTPSs in this study. Blind docking using CB-Dock2 can be used to study the binding properties and molecular mechanisms between protein and substrate, to reveal key residues that are functionally relevant in the binding pocket(Alvarez et al., 2021; Chen et al., 2023; Liu et al., 2022). Based on the structural data, it has been observed that the variable arrangement of the *γ, β,* and *α* domains in PdiTPSs (Fig. 2) is an important strategy for expanding and diversifying diterpene synthases. Their combination, presence, and absence constitute the structural chemistry of diterpene synthases (Faylo et al., 2021; Koksal et al., 2011; Zhou et al., 2012).

In addition, the correlation analysis showed that the similarity of overall structures increased with the similarity of overall sequences (PCC = 0.78, *p* < 0.001). The N-terminal subsequence showed high correlation with the overall sequence and moderate correlation with the TM score of the overall structure (PCC = 0.69, *p* < 0.001). Conversely, the C-terminal subsequence differed significantly from the N-terminal and overall sequences, and exhibited weak correlation with the overall structure (PCC = 0.33, *p* < 0.001). When analyzing the correlation between residues surrounding the substrate and the overall structure, the trend was completely opposite (Supplementary Table S5). The average correlation between the C-terminal subsequence (0.54) and residues around the substrate was higher than that between the N-terminal subsequence (0.37). Combining the N- and C-terminal subsequence as the NC-terminal subsequence increased the average correlation with residues around the substrate to 0.58, which is understandable because both N- and C-terminal subsequences contain residues around the substrate. Furthermore, there was no correlation between residues around the substrate and the topology of the overall structure, as evidenced by the TM score distribution of residues around the substrate, which was significantly lower than that of the overall structure (Figs. 4C and 4D). This suggested that the overall structure of the PdiTPSs might has a different folding mechanism from the topology formed by the residues surrounding the substrate.

To evaluate the relationships between sequence similarity, TM score of protein structures, and products, the impact of sequence similarity and TM score on products was indirectly reflected by measuring the strength of their correlation. The similarity between products was calculated, and the final correlation coefficients were summarized in Table 1. Surprisingly, the overall sequence of PdiTPSs had the highest correlation with the product, while the residues around the substrate and the overall topology had a weak correlation with the product. The N-terminal showed stronger correlation with products than that of the C-terminal, which was consistent with our previous grouping of PdiTPSs and products based on SSN mapping.

**Table 1.** PCC for PdiTPSs sequences, structures, and products

| *Content* | *Pearson correlation coefficient* |
| --- | --- |
| 4Å : Product | 0.38 |
| 6Å : Product | 0.4 |
| 8Å : Product | 0.39 |
| 10Å : Product | 0.4 |
| overallstructure : Product | 0.36 |
| Csequence : Product | 0.35 |
| Nsequence : Product | 0.54 |
| NCsequence : Product | 0.54 |
| overallsequence : Product | 0.55 |

Note: All $p$ values $< 0.001$ in the table's statistical results.

To further validate the impact of the correlation between product and sequence similarity on product grouping, the similarity mapping between product and conserved motifs in PdiTPSs was analyzed by MEME. The four signature motifs (LHS, PNV, FERLW, and PIX) located in l different long motifs (Fig. 5) had been identified. The similarity of these motifs to product was then correlated with product similarity. The correlation between the similarity of motifs 1 (PNV and FERLW) and product was 0.51, while the correlation for motif 3 (PIX) and motif 2 (LHS) were 0.32 and 0.36, respectively. Motifs with higher product correlation have more mutated residues in non-functional motifs, and conversely, possess fewer mutations. This correlation might indirectly reflect the level of differentiation between enzyme motifs, where motifs with high product correlation likely represented functional domains of the enzyme, while positions with low product correlation might contain product-specific motifs.

**Fig. 5.**
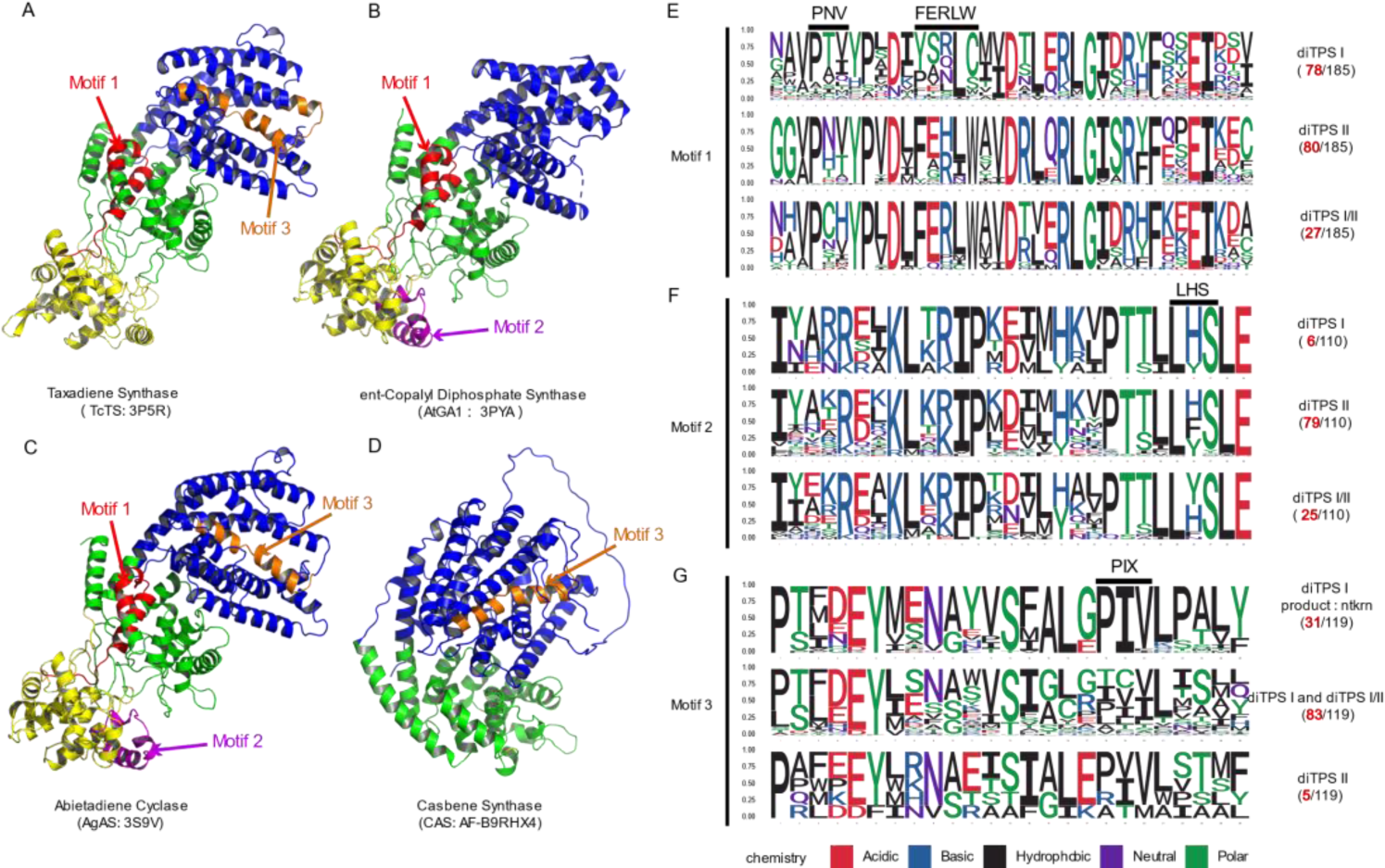
The positions of validated functional motifs in protein structures, along with the number of motifs and specific residues near the motifs in the three classes of PdiTPSs. A) The crystal structure of Taxadiene synthase (diTPS I); B) The crystal structure of *ent*-copalyl diphosphate synthase (diTPS II); C) The crystal structure of Abietadiene cyclase (diTPS I/II); D) The crystal structure of Casbene synthase (diTPS I). E) Seqlogo of the PNV and FERLW motifs and their neighboring residues; F) Seqlogo of the LHS motif and its neighboring residues; G) Seqlogo of the PIX motif and its neighboring residues. The black numbers represent the total number of sequences retrieved by MEME for the PNV, FERLW, LHS, and PIX motifs, while the red numbers represent the number of occurrences of each motif in the three classes of enzymes.

### 2.8 Aromatic residues around PdiTPSs substrates determine substrate type

Generally, the probability of catalytic associated residues appearing elsewhere in the sequence should be lower than that of appearing at the catalytic center. The statistical analysis revealed that aromatic amino acids around substrates with different sizes were the most frequently observed in PdiTPSs (Figs. 6A and 6B). This might be due to the fact that the aromatic rings in aromatic amino acids could donate electrons to stabilize the carbocation intermediates in the catalytic process of PdiTPSs, making it easier for the substrates to be converted into products. Specifically, tryptophan (W) was more likely to occur around the linear substrate of GGPP, while phenylalanine (F) and tyrosine (Y) were more likely to present around the substrate of the initial cyclization intermediate of PdiTPSs (Figs. 6C and 6D).

**Fig. 6.**
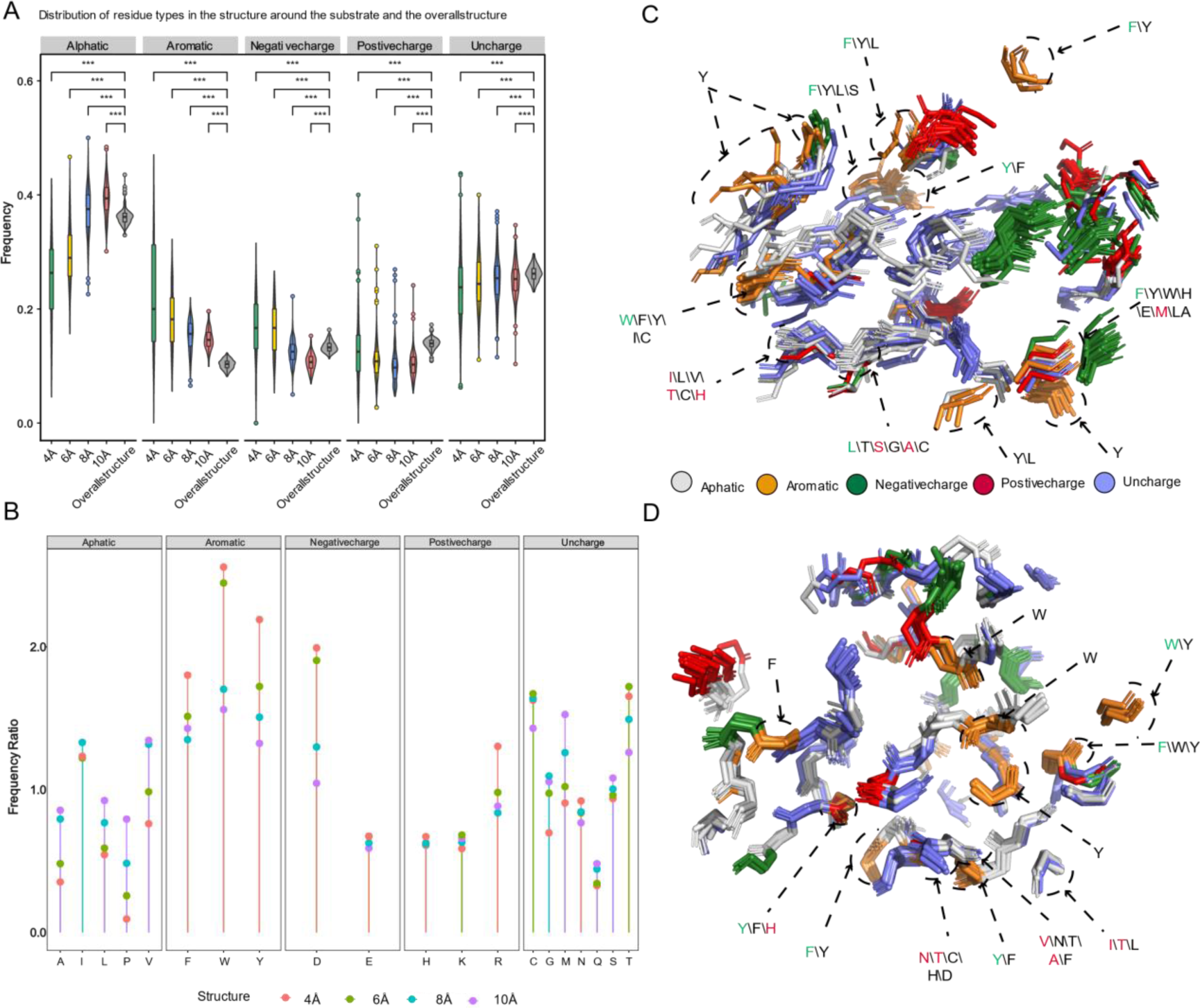
Preference of 5 types of amino acids in the residues around the substrate and their relative spatial position in the structure. A) Frequency distribution of five types of amino acids (aliphatic, aromatic, negatively charged polar, positively charged polar, and uncharged) within 4 Å, 6 Å, 8 Å, and 10 Å from the substrate compared to their overall frequency within the protein structure. B). Comparison of the frequency of 20 amino acids within the protein structure to their frequency within 4 Å, 6 Å, 8 Å, and 10 Å from the substrate. C) Superimposition of substrate-proximal residues (within 6 Å) of 15 diterpene synthases producing different skeletons, as well as those of diTPS I and diTPS I/II that have undergone mutagenesis studies. D) Superimposition of substrate-proximal residues (within 8 Å) of diterpene synthases producing different intermediates, as well as those of diTPS II and diTPS I/II that have undergone mutagenesis studies. In C and D, aromatic residues and residues that have been shown to affect product outcome in mutagenesis studies are highlighted. Green fonts indicate the residue with the highest frequency in that position, and red fonts indicate mutated residues (also with the highest frequency). Asterisks indicate statistical significance (*: p < 0.05, **: p < 0.01, ***: p < 0.001), and “NS” indicates no statistical difference.

Studies on the impact of aromatic residues on the function of diterpene synthases have only indicated that the active sites of Class I and II enzymes both contain at least 2-3 aromatic residues, which guide the intermediate involved in the reaction through spatial constraints and cation-π interactions. Moreover, the number of aromatic residues in the active site can be used to predict the promiscuity of the enzyme (Zhang et al., 2020). However, there is no report explaining the selectivity of substrate structure exhibited by the observed types of aromatic residues.

### 2.9 Residues within 8 Å of substrate have more impact on products

It had been analyzed for the residues surrounding the substrate of PdiTPSs that generated different diterpene scaffolds and intermediates. The residues whose mutation had been experimentally demonstrated to affect product formation were also examined. It had been found that the physicochemical properties and spatial orientations of these residues surrounding the substrate were highly conserved. Some frequently occurring residues, such as arginine (R), cysteine (C), tryptophan (W), aspartic acid (D), isoleucine (I), serine (S), threonine (T), valine (V), phenylalanine (F), tyrosine (Y), and methionine (M) (Fig. 6B), were also conserved in their spatial positions (Figs. 6C and 6D). Except for arginine (R), cysteine (C), phenylalanine (F), and tryptophan (W), the effects of the other residues on enzyme function and product had been demonstrated by mutagenesis experiments. For example, the PdiTPSs OsKSL5i: I664T and OsKSL5i: I718V from rice (*Oryza sativa*) specifically produced *ent*-pimara-8(14),15-diene and *ent*-isokaur-15-ene, respectively (Xu et al., 2007); Six PdiTPSs from *Tripterygium wilfordii* independently evolved new functions by mutating specific residues, including TwKSL1v2: M607\T638A, TwKSL3: M608\I639, TwKSL2: A608\I639, TwCPS3: I115\N327\V328\H268, TwCPS5: T115\A327\T326, and TwCPS6: Y265 (Tu et al., 2022). Moreover, mutation of glutamic acid (E) at position 690 to arginine (R), phenylalanine (F), lysine (K), proline (P), or aspartic acid (D) or mutation of serine (S) at position 721 to valine (V) in SmMDS resulted in product loss (Tong et al., 2022), respectively.

It had been also observed that some low-frequency residues, such as alanine (A) and histidine (H), contributed to product specificity. For example, the AgAS: A723S mutant of abietadiene synthase specifically produced pimaradienes, and the H268 residue in TwCPS3 also contributed to product specificity (Tu et al., 2022; Wilderman and Peters, 2007). Additionally, these low-frequency residues around the substrate often appeared in spatially conserved but physicochemically lowly conserved positions (Figs. 6C and 6D). Mutated residues affecting the product mainly occurred within 6 Å of the substrate in PdiTPS I and within 6 Å to 8 Å of that in PdiTPS II. (Figs. 6C and 6D). For specific residue shapes and maps within 4 Å to 8 Å of representative PdiTPS I and PdiTPS I/II that produced SK1-SK15 skeleton types, as well as that of PdiTPS II, please referred to the supplementary data (Figs. S3-S21).

The previous analysis of residues around the substrate can provide clues to the importance of residues, but the effect of the residue on the product outcome still needs to consider factors such as spatial position and interaction with adjacent residues around the substrate. Nevertheless, these valuable experimental results also provide important evidence for the important position pattern of residues located within 8 Å of the substrate in PdiTPSs proteins that determines the product outcome. Our study is only a starting point, and in the future, we can obtain a new structural motif of conserved residues in PdiTPSs by examining and modifying structure alignment to derive sequence-order independent structural site motifs (Sankar and Chandra, 2022; Sankar et al., 2022). The study of these structural motifs may be a window for us to view the spectacular chemical diversity of PdiTPSs.

## 3. Conclusion

In this study, we have compiled a dataset of plant-derived diterpene synthase with sequences and structural information, which is the largest annotated collection of PdiTPSs to date. It covers mosses, ferns, gymnosperms, and angiosperms. The dataset can be applied for studies of sequence and structure analysis, as well as serving as a library of enzyme components for combinatorial biology experiments to obtain non-natural diterpene products and a wider range of diterpene derivatives (Jia et al., 2019). The dataset also offers a method for grouping diterpene products based on their backbone, which simplifies the analysis and comparison of enzyme function. This method may be useful for researchers studying diterpene synthesis in plants. It has been discovered for the strong correlation between N-terminal subsequence and overall sequence, significant sequence differences between N- and C-terminal subsequences from this data. Structural conservation increases with increasing similarity between the N-terminal sequence and the overall sequence. Moreover, an independent topological structure exists between the residues around the substrate and the overall structure. Quantitative analysis showed that sequence similarity has a greater impact on product distribution than structural similarity or similarity in the topological structure of residues around the substrate. However, some residues within 8 Å of the substrate have a profound effect on product. Furthermore, it has been found that tryptophan (W) is specific to the binding cavity of PdiTPSs that recognize linear substrate GGPP, while phenylalanine (F) and tyrosine (Y) are specific to the binding cavity of PdiTPSs that recognize cyclic substrates. Our exploration of the features of PdiTPSs and their mapping to product outcomes can be applied to enzyme identification, engineering, and product prediction research. In summary, our curated dataset will help to understand the sequence and structural space of PdiTPSs and provide a starting point for the specificity analysis of enzyme-controlled product distribution.

## 4. Methods

### 4.1 Collect characterized PdiTPSs

To find potentially characterized PdiTPSs, it has been manually searched for evidence of experimental characterization of diterpenes from the literature up to May 2022 and collected their corresponding GenBank accession numbers (NCBI). HMM of the N-terminal domain (PF01397) and C-terminal domain (PF03936) sequences of terpene synthases were downloaded from the Pfam database. Hmmsearch v.3.1.2 was used to search for the N- and C-terminal sequences in each PdiTPSs sequence. When multiple N- or C-terminal sequences were identified, the result with the lowest e-value was retained.

### 4.2 Construction of sequence similarity networks

All-versus-all pairwise local sequence alignments were performed using SSNpipe v.1.0.0 for PdiTPSs C, N, NC domains, and overall sequences (Viborg et al., 2019). The BLAST result files were searched with E-value thresholds ranging from 10^-5^ to 10^-140^ at 5 log unit intervals. The network files were visualized using Cytoscape (http://www.cytoscape.org/).

### 4.3 Phylogenetic analysis and visualization

The protein sequences were aligned using MAFFT v.7.310 (Katoh et al., 2002) with the following parameters: maxiterate 1000 –localpair –thread 30--maxiterate 1000 --genafpair --thread 30. --maxiterate 1000 --globalpair --thread 30. Manual inspection was performed to ensure proper alignment of known motifs such as the DDXXD and DXDD motif. Phylogenetic tree was inferred using IQTree v.2.0.3 (Nguyen et al., 2015) with the following parameters: -s Mafft_Sequence -m JTT+F+R7 (full-length)/JTT+F+R5 (C-terminal and N-terminal subsequences) –B 1000 -nt AUTO. Levopimaradiene synthase from the hornwort *Phaeoceros carolinianus* was specified as the outgroup, and the resulting tree was visualized by Chiplot (https://www.chiplot.online/tvbot.html).

### 4.4 Retrieve and visualize sequence motifs

199 PdiTPSs amino acid sequences were submitted to the MEME online tool (https://meme-suite.org/meme/tools/meme) to identify motifs. Based on the known functional widths, the number of motifs was set to 20, and the minimum width of the motifs was set to 3. Other parameters were set to default values.

### 4.5 Perform calculations of similarity and correlation between sequences, structures, and small molecules

The similarities between C-terminal, N-terminal, and overall sequences were compared by using TBtools (Chen et al., 2020) Protein Pairwise Similarity Matrix. TMalign (Zhang and Skolnick, 2005) was used to compare the topology similarity between the residues around the substrate and the overall structure, with the command “TMalign Pdb-A Pdb-B -outfmt 2”, and the structure similarity was expressed as the TM score. The Dice Similarity Coefficient (DSC) (Willett et al., 1998) was used to measure chemical similarity in extended connectivity fingerprint (ECFP/Morgan Fingerprint, radius 2). For each pair of PdiTPSs, their corresponding products were arranged and combined, and the similarity score was calculated using the RDKIT similarity matrix. For PdiTPSs that produce only one product, there is only one product pair and thus only one similarity value. However, for those that produce multiple products, several product pairs were obtained, and the average of their similarity scores was used to represent the product similarity values. The pearson correlation coefficient (PPC) was calculated using ggstatsplot in R. The correlation analysis involved the following factors: the overall sequences of 199 PdiTPSs, the sequence similarity of C-terminal, N-terminal, and NC terminal subsequences, as well as the structure formed by residues surrounding the substrate and the overall structure.

### 4.6 Perform structure prediction, molecular docking and visualization

153 structures of PdiTPSs were downloaded from the AlphaFold database (https://alphafold.com/), and the rest were predicted for their 3D structures using local AlphaFold2. The obtained structures were docked with substrates using the CB-Dock2 molecular docking program (https://cadd.labshare.cn/cb-dock2). Structures were docked with substrates using the CB-Dock2 molecular docking program (https://cadd.labshare.cn/cb-dock2). The docking postures outputted by CB-Dock2 were compared with the crystal structures of PdiTPSs, and the complex closest to the crystal structure was selected as the final docking result. The amino acids within 4-10Å of the substrate were selected using the command “select AA, byres all within 4 of sele” in PyMOL (version 2.0) for further analysis. The diagrams were made with PyMOL.

### 4.7 Calculate amino acid frequencies

Software iLearnPlus v.1.0.1 (Chen et al., 2021) was used to extract the AAC and GAAC features from the residues surrounding the substrates of different sizes and the overall sequences. The resulting features included the frequencies of the 20 amino acids and their categorization based on 5 physicochemical properties. The residue preferential values were then calculated the ratio of frequency of each residue around the substrate to its frequency in the overall sequence.

## Data availability

Data will be made available on request.

## Acknowledgments

This work was supported by the National Natural Science Foundation of China (31760189).

