## Supplementary material for "Exploration of an enzyme-product mapping approach for plant-derived diterpene synthases": Figures and Table S5

**Supplementary Figures**

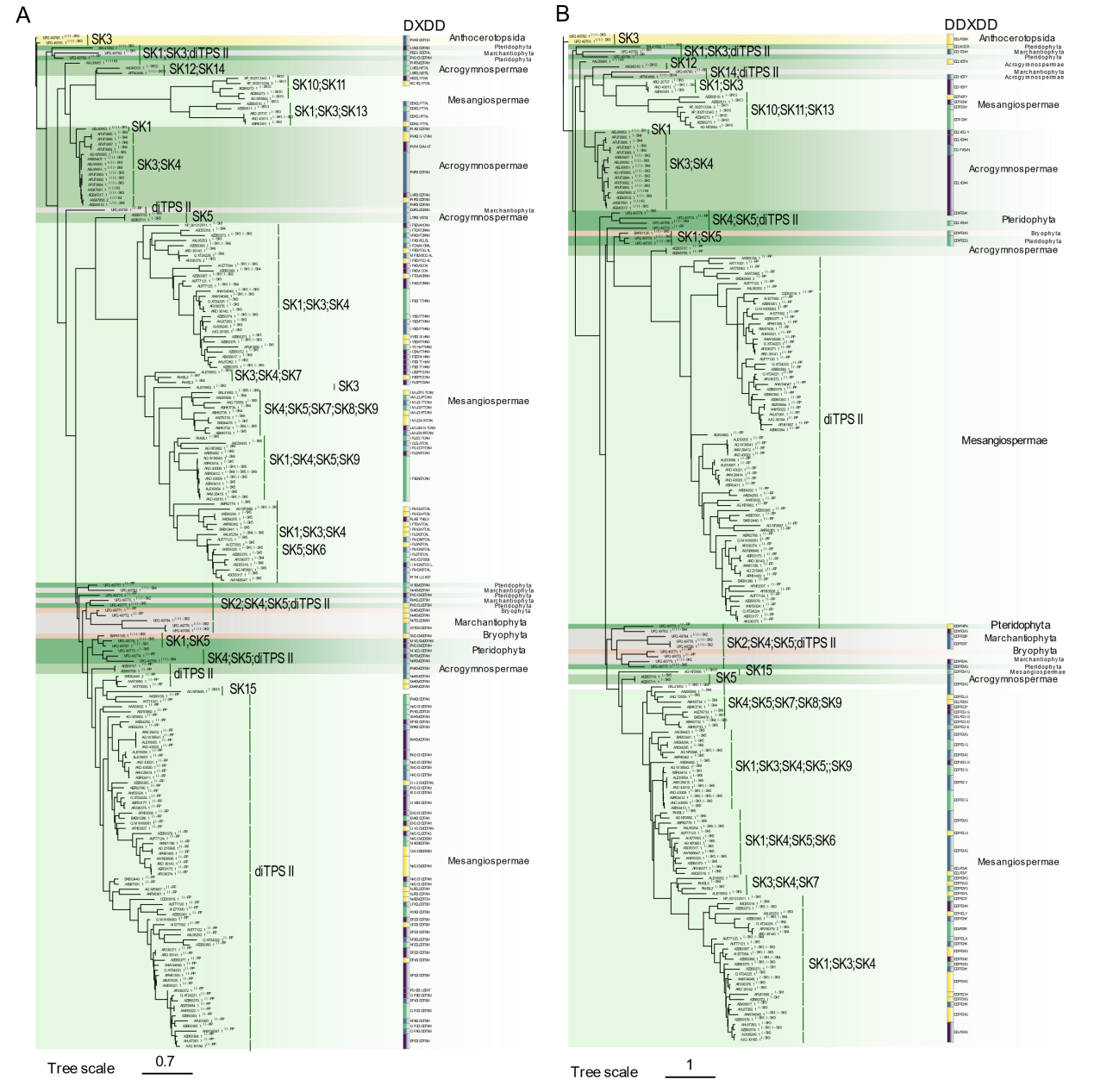

**Fig. S1**. The relationship between the phylogeny of the N-terminal and C-terminal domains of PdiTPSs the motifs related to product skeleton and function, and the classification level of enzyme sources. The phylogenetic tree indicates the ID, product type, and product skeleton classification of each PdiTPSs. A) N-terminal domain, DXDD is the signature motif of the N-terminal domain; B) C-terminal domain, DDXXD motif is the signature motif of the C-terminal domain. The extended motifs of these two motifs are shown here.

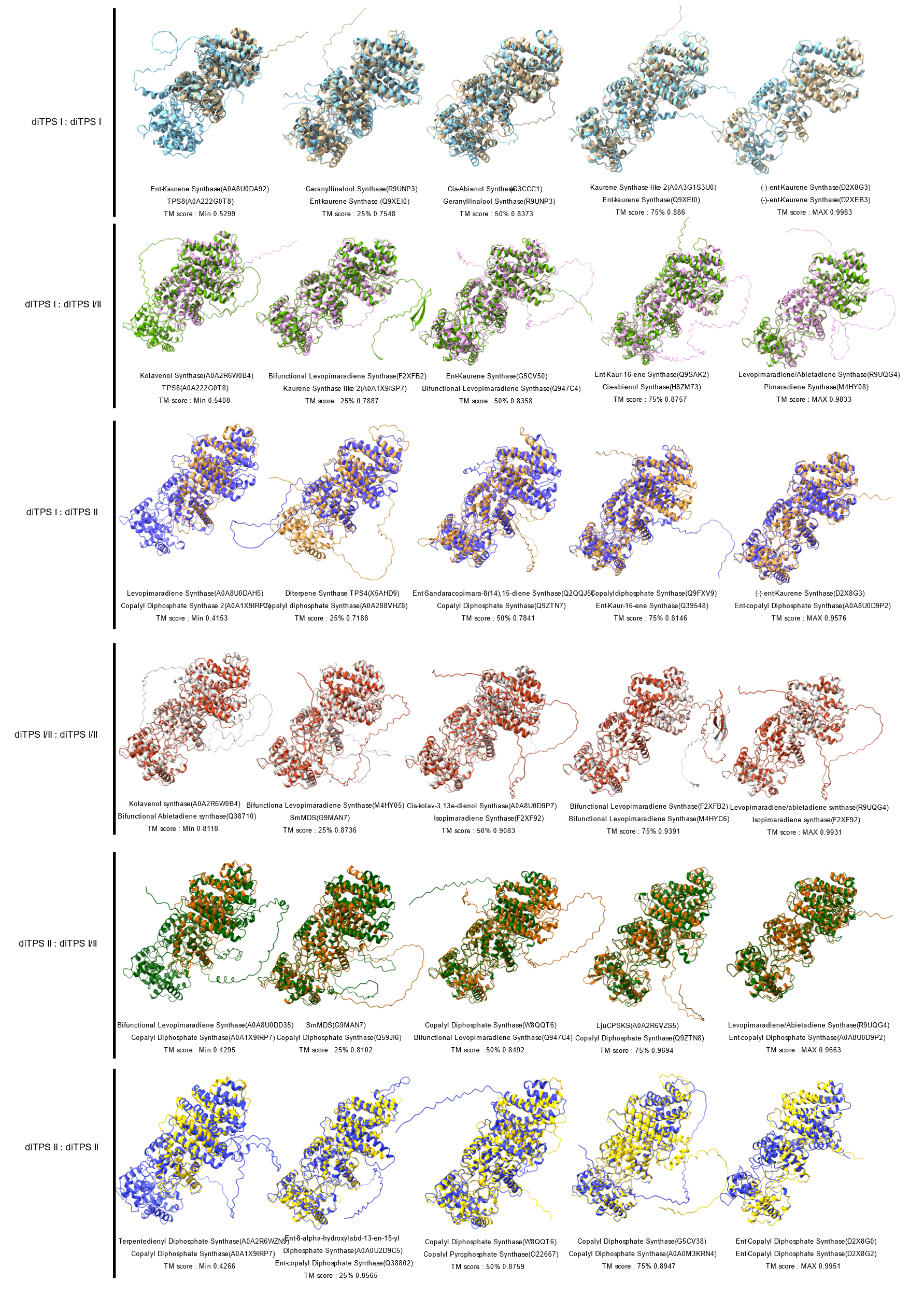

**Fig. S2**. Representative structural superpositions of the minimum, 25% (Q1), 50% (Q2), 75% (Q3), and maximum observed values of the distribution of topological similarity between PdiTPS I, PdiTPS II, and PdiTPS I/II.

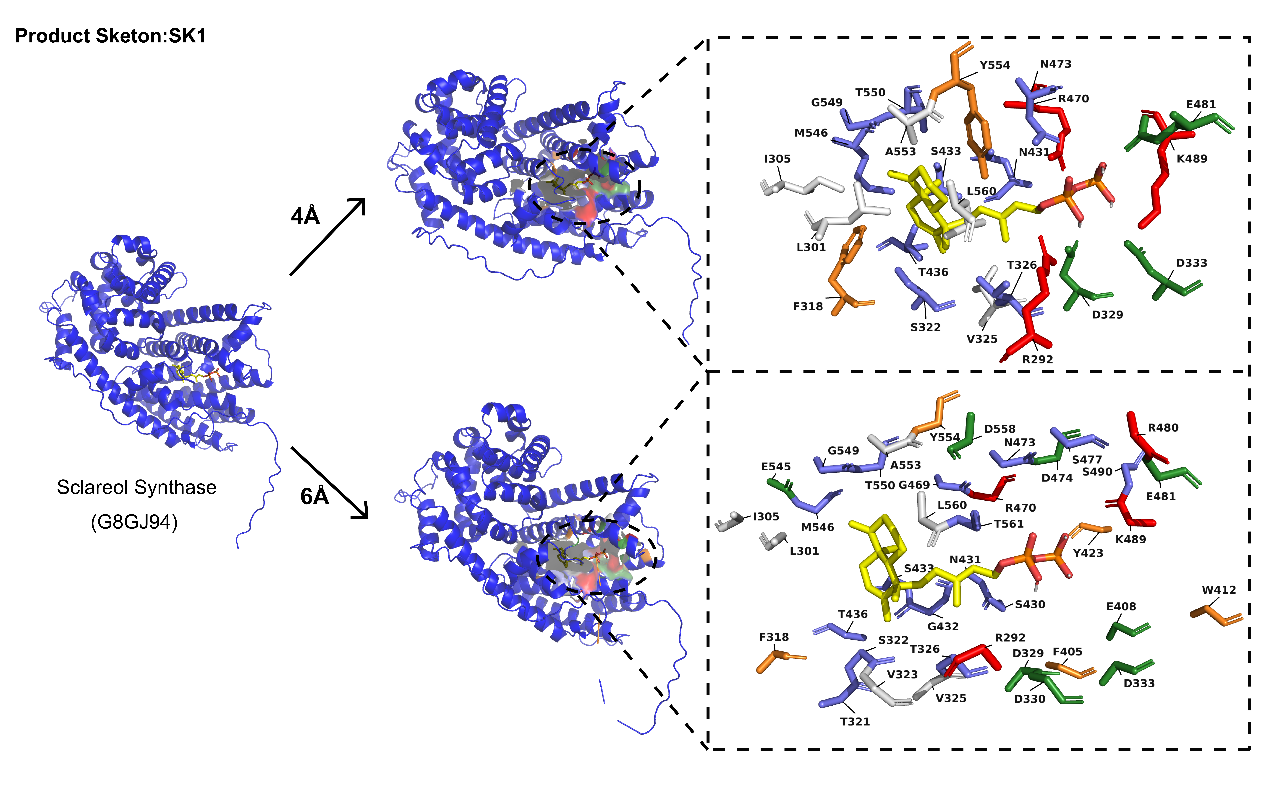

**Fig. S3**. Specific residue structures and topology formed by residues at 4Å and 6Å radial distances from the substrate of PdiTPSs producing the SK1 skeleton type.

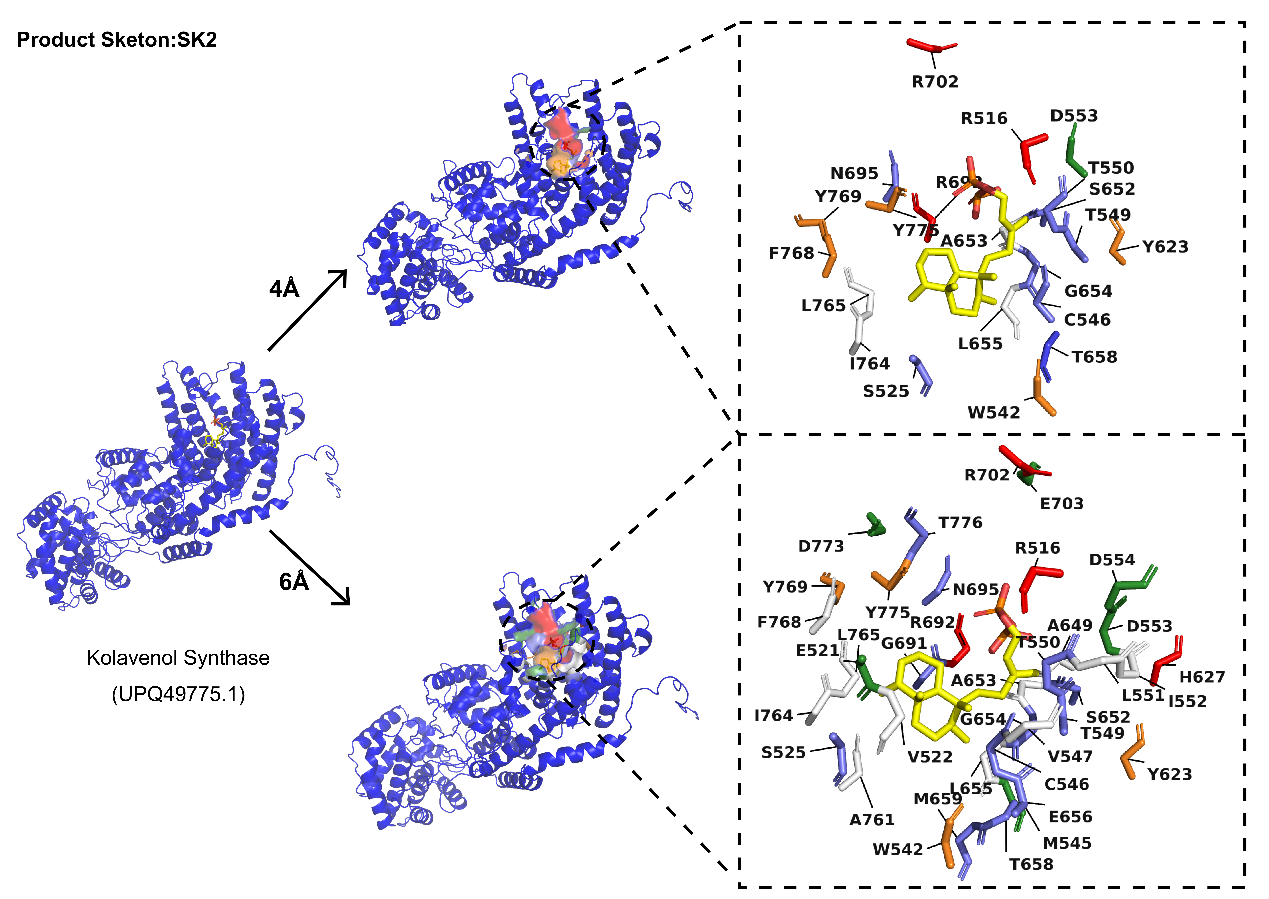

**Fig. S4**. Specific residue structures and topology formed by residues at 4Å and 6Å radial distances from the substrate of PdiTPSs producing the SK2 skeleton type.

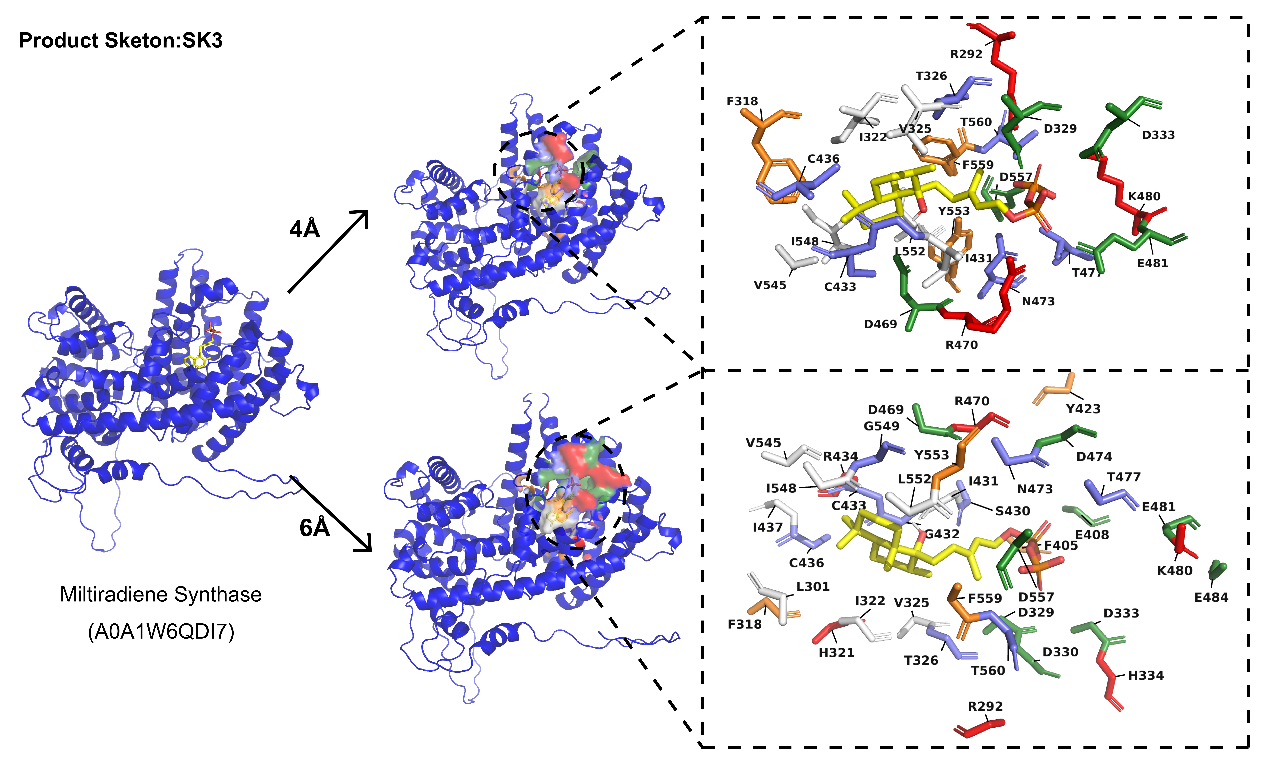

**Fig. S5**. Specific residue structures and topology formed by residues at 4Å and 6Å radial distances from the substrate of PdiTPSs producing the SK3 skeleton type.

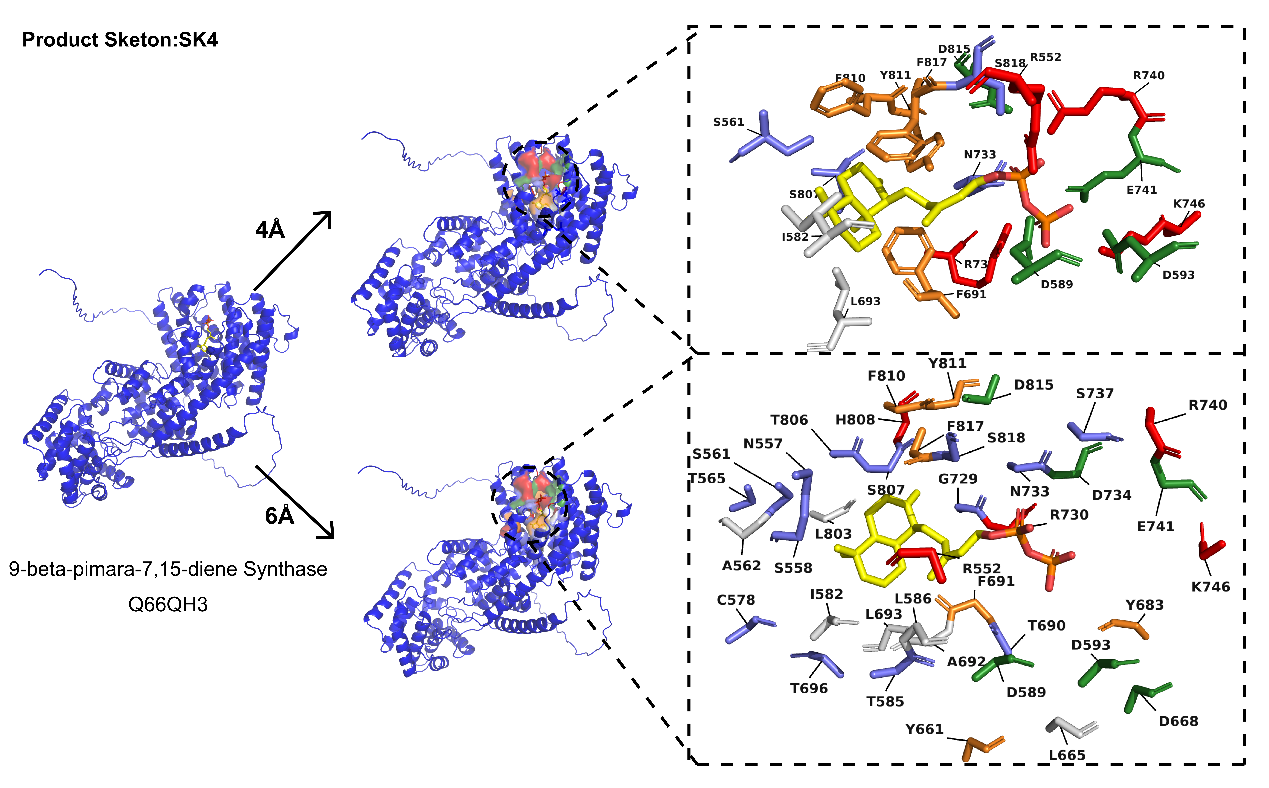

**Fig. S6**. Specific residue structures and topology formed by residues at 4Å and 6Å radial distances from the substrate of PdiTPSs producing the SK4 skeleton type.

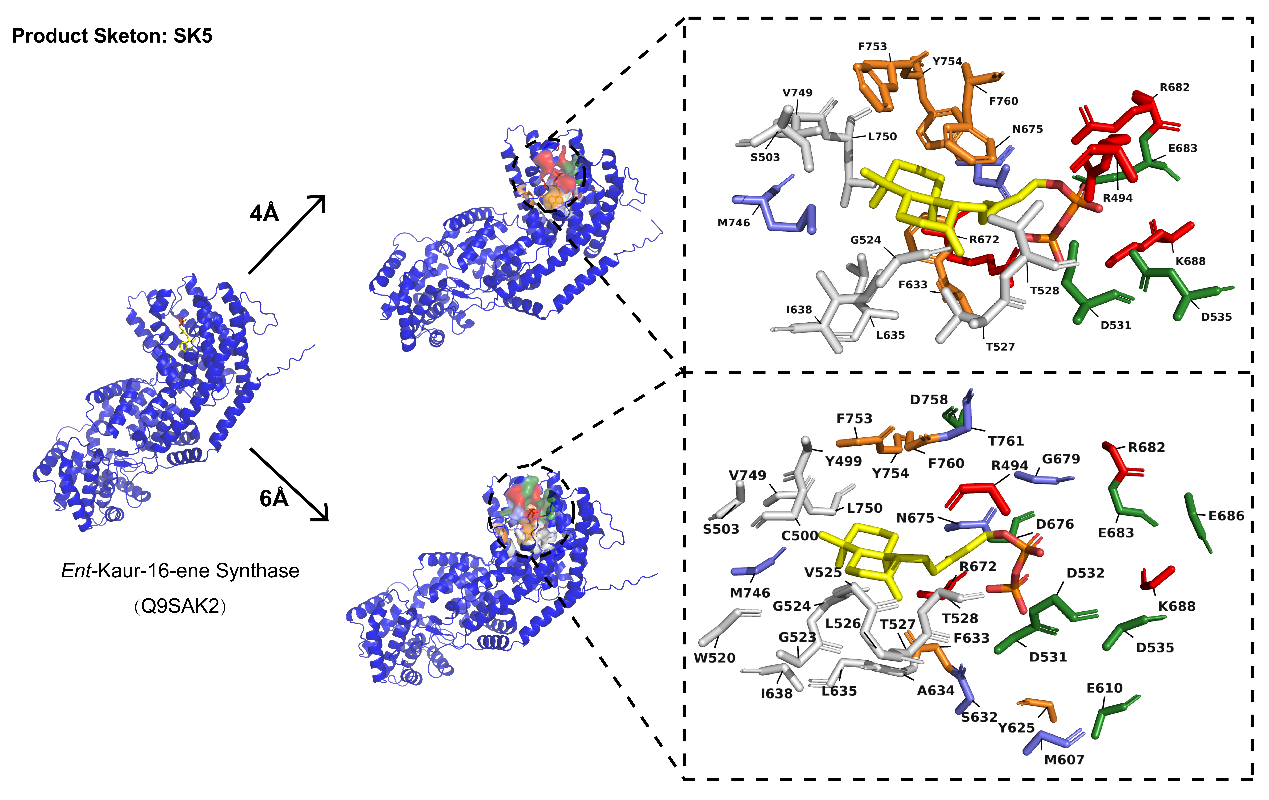

**Fig. S7**. Specific residue structures and topology formed by residues at 4Å and 6Å radial distances from the substrate of PdiTPSs producing the SK5 skeleton type.

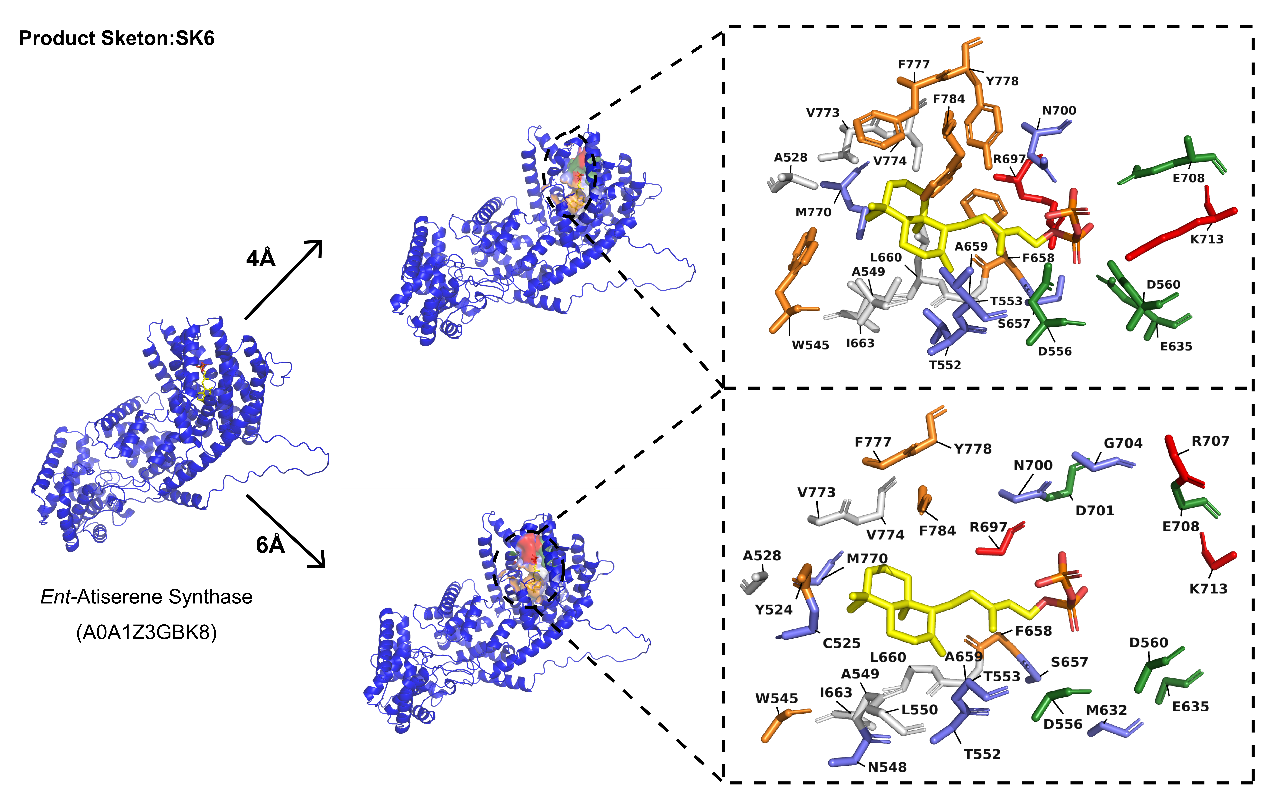

**Fig. S8**. Specific residue structures and topology formed by residues at 4Å and 6Å radial distances from the substrate of PdiTPSs producing the SK6 skeleton type.

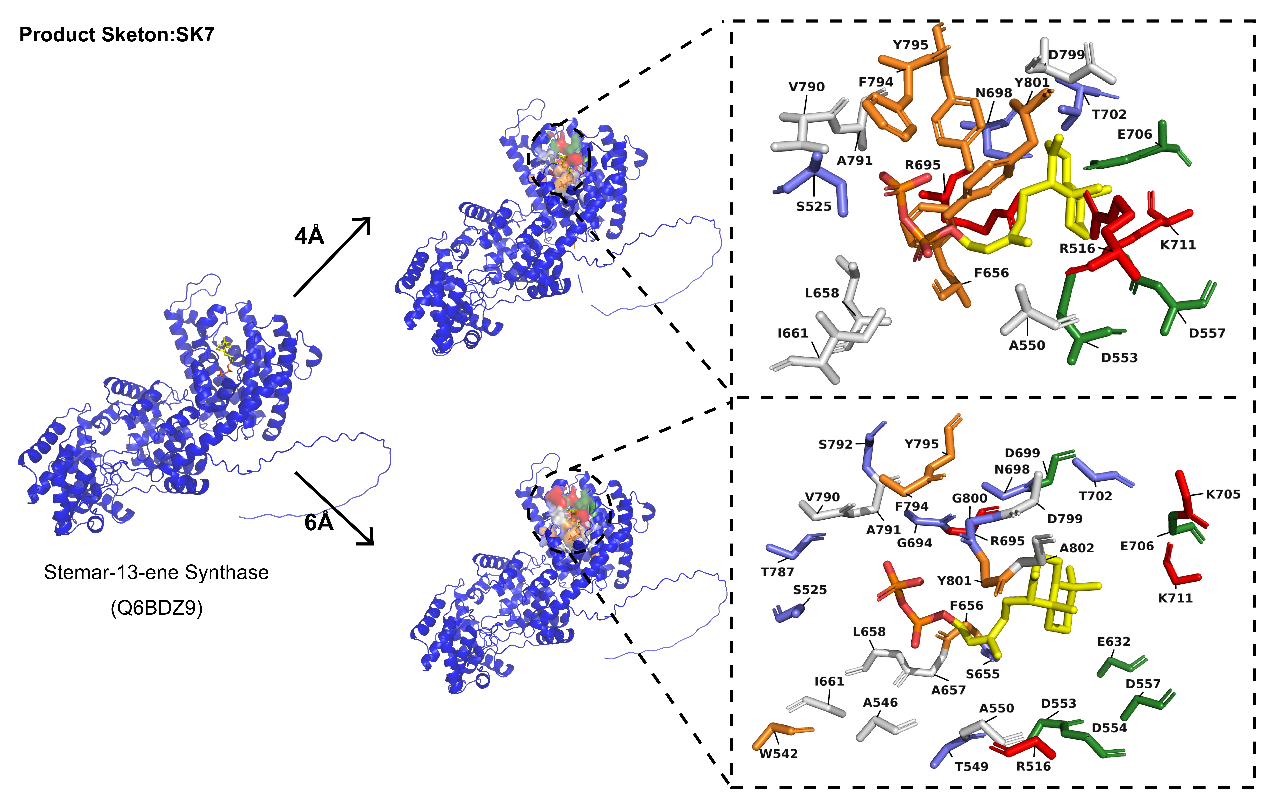

**Fig. S9**. Specific residue structures and topology formed by residues at 4Å and 6Å radial distances from the substrate of PdiTPSs producing the SK7 skeleton type.

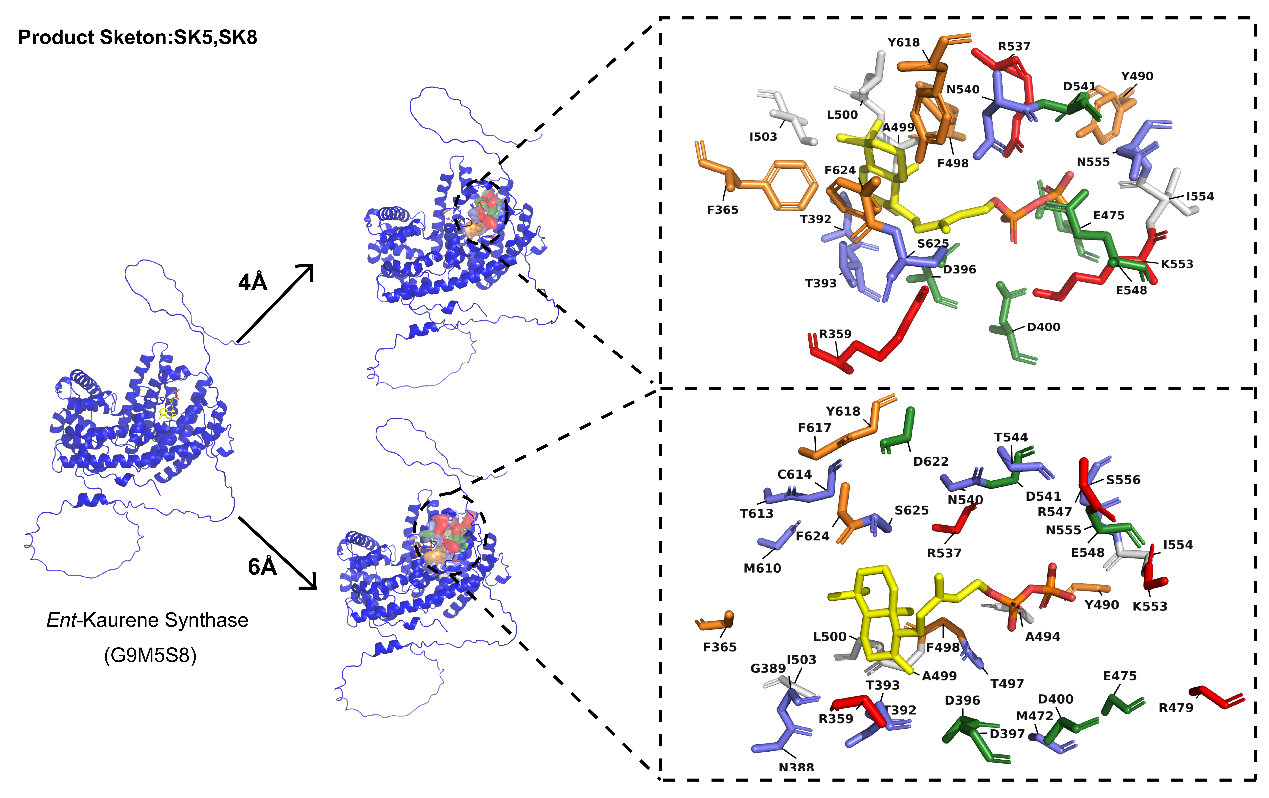

**Fig. S10**. Specific residue structures and topology formed by residues at 4Å and 6Å radial distances from the substrate of PdiTPSs producing the SK8 skeleton type.

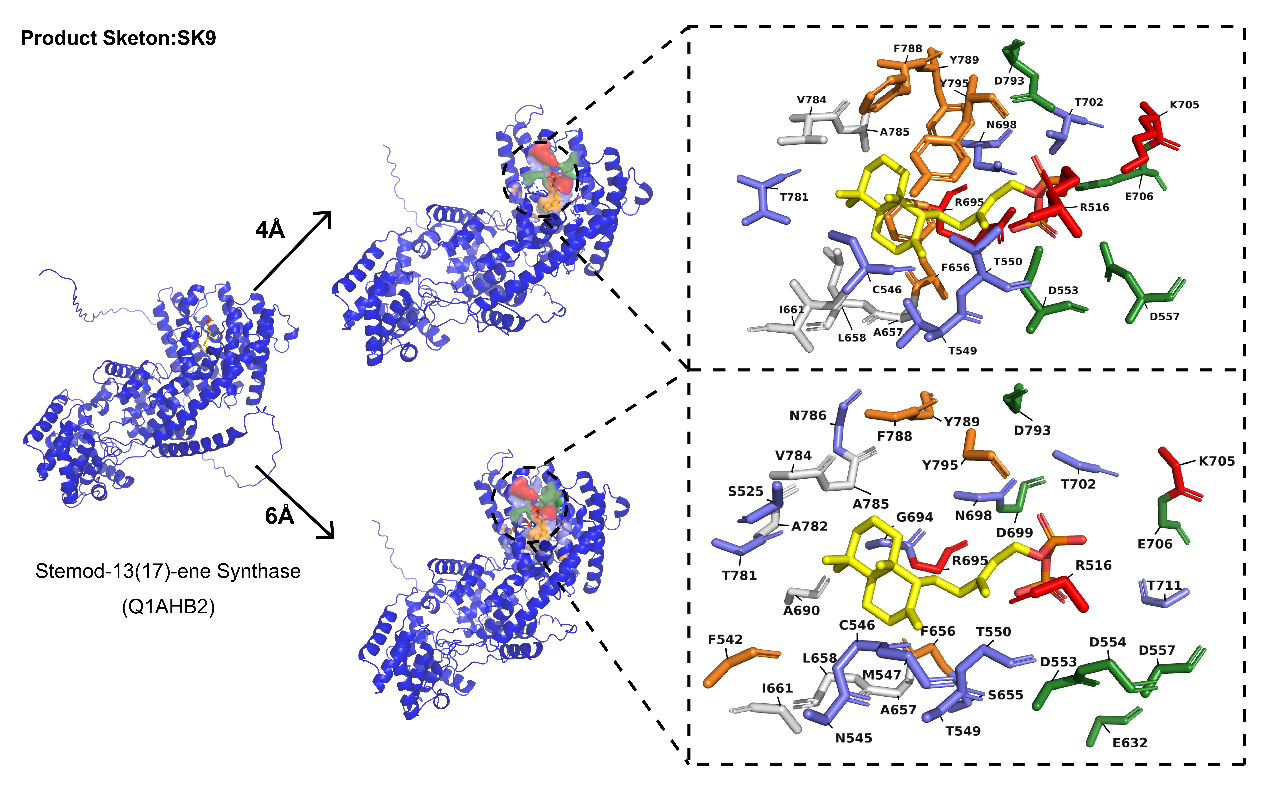

**Fig. S11**. Specific residue structures and topology formed by residues at 4Å and 6Å radial distances from the substrate of PdiTPSs producing the SK9 skeleton type.

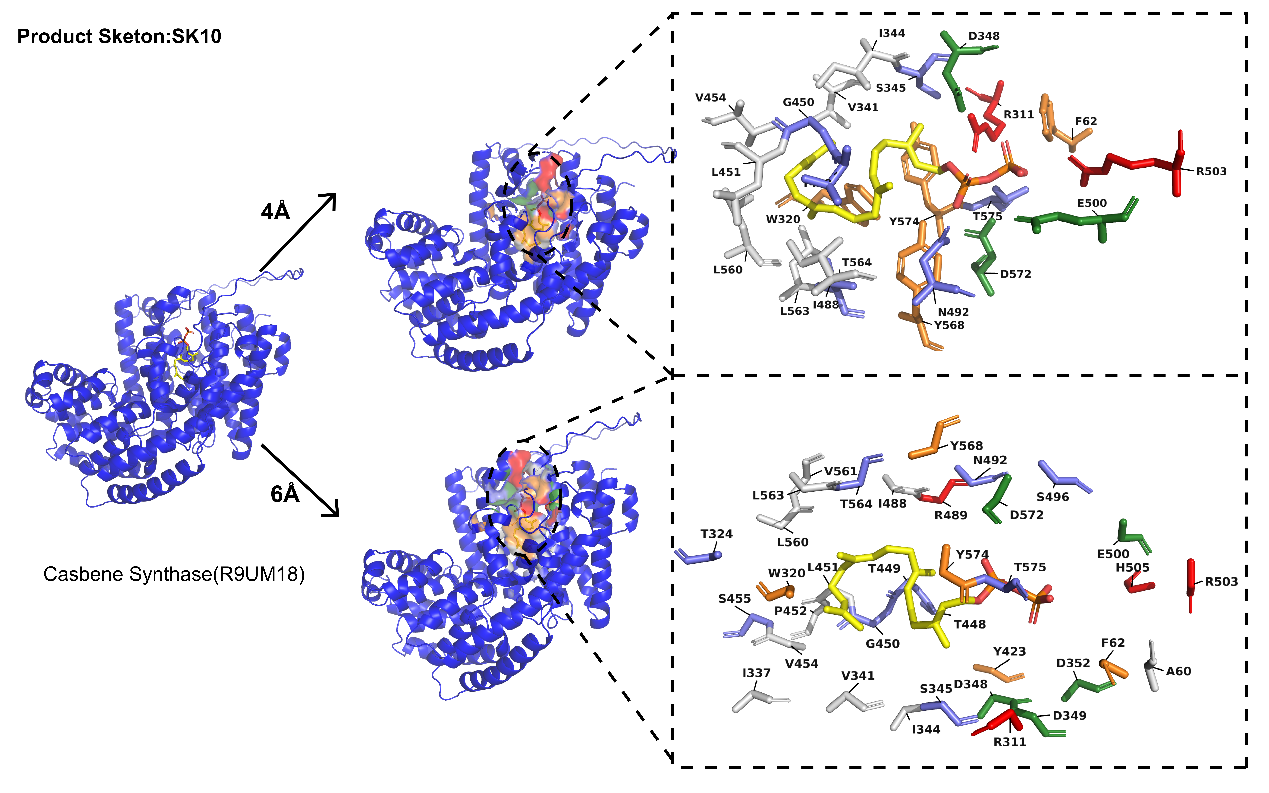

**Fig. S12**. Specific residue structures and topology formed by residues at 4Å and 6Å radial distances from the substrate of PdiTPSs producing the SK10 skeleton type.

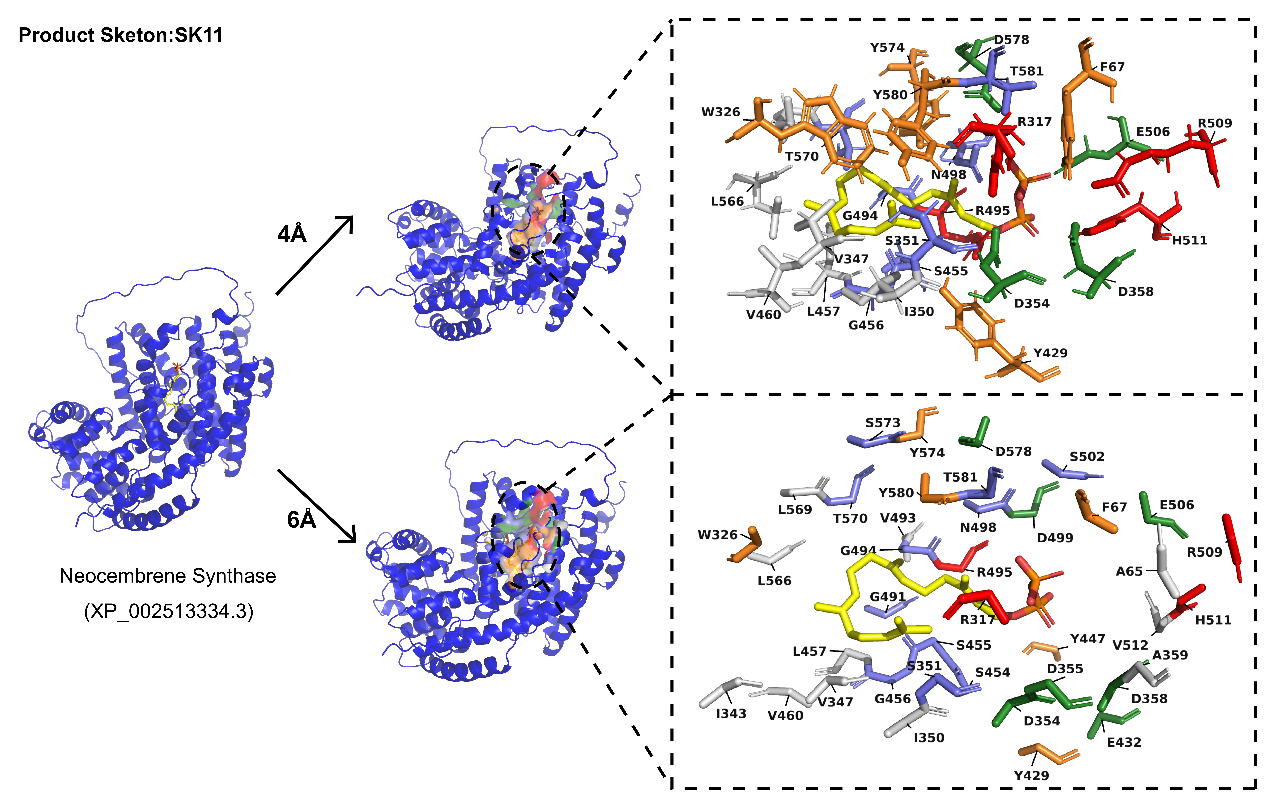

**Fig. S13**. Specific residue structures and topology formed by residues at 4Å and 6Å radial distances from the substrate of PdiTPSs producing the SK11 skeleton type.

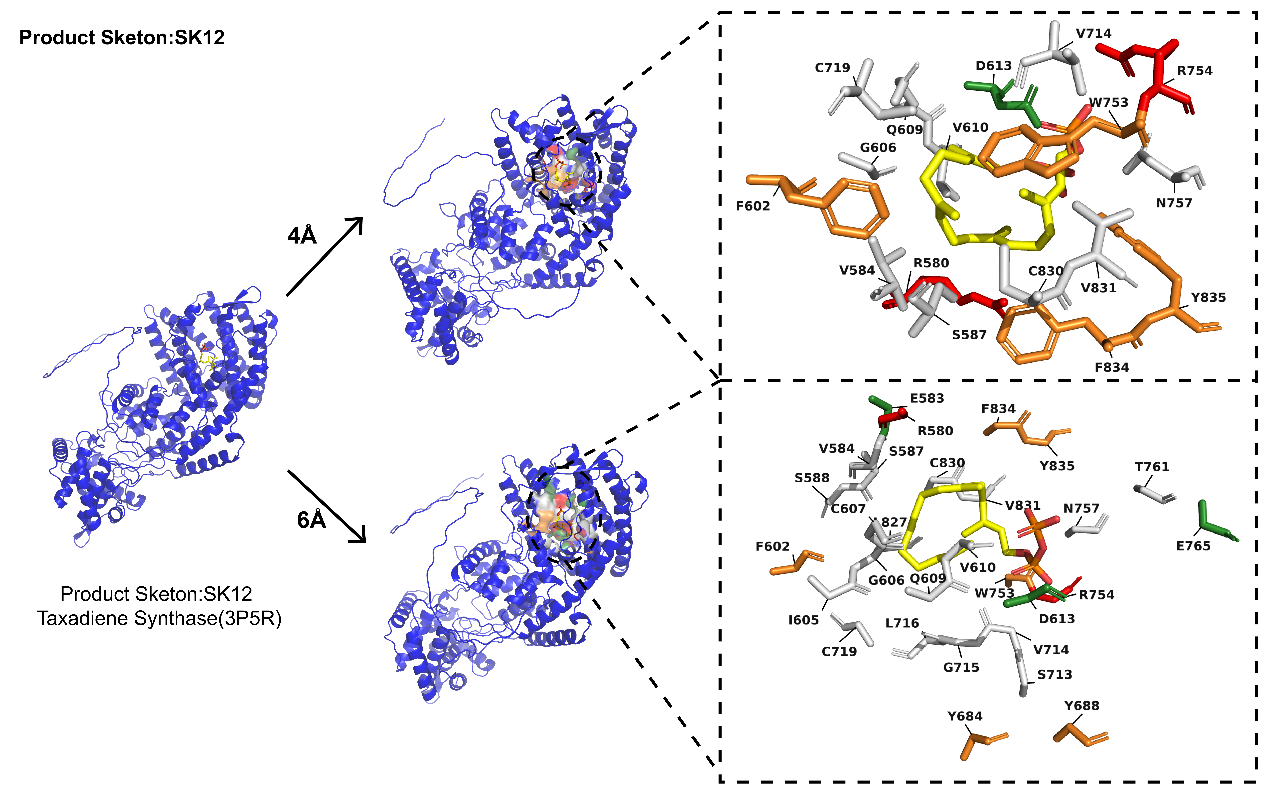

**Fig. S14**. Specific residue structures and topology formed by residues at 4Å and 6Å radial distances from the substrate of PdiTPSs producing the SK12 skeleton type.

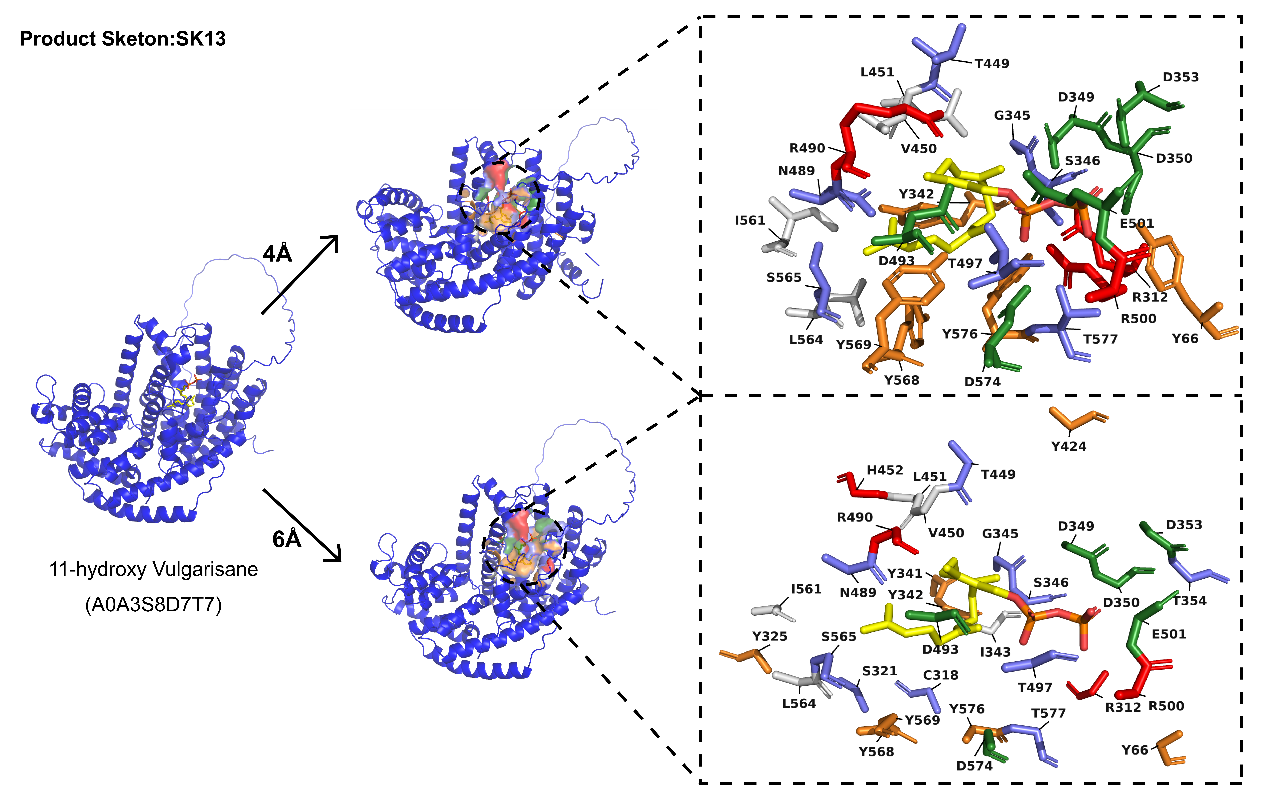

**Fig. S15**. Specific residue structures and topology formed by residues at 4Å and 6Å radial distances from the substrate of PdiTPSs producing the SK13 skeleton type.

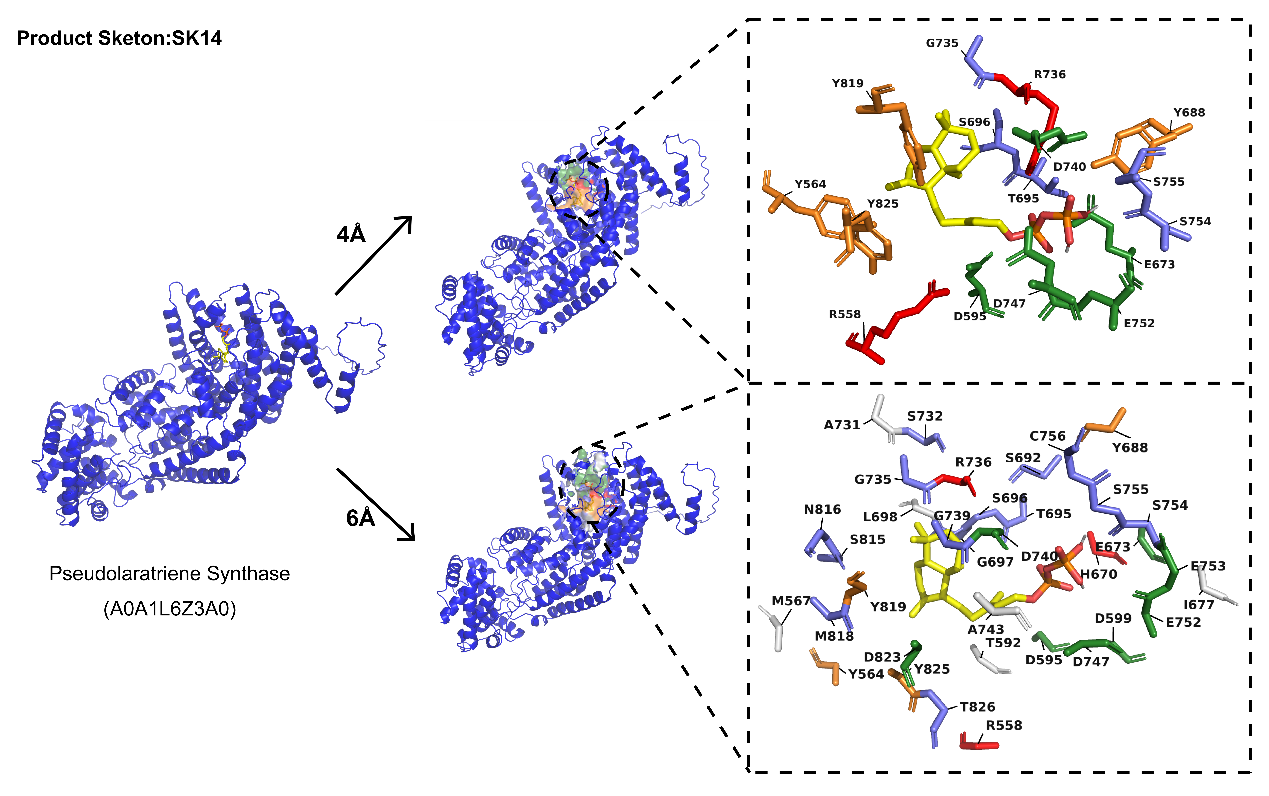

**Fig. S16**. Specific residue structures and topology formed by residues at 4Å and 6Å radial distances from the substrate of PdiTPSs producing the SK14 skeleton type.

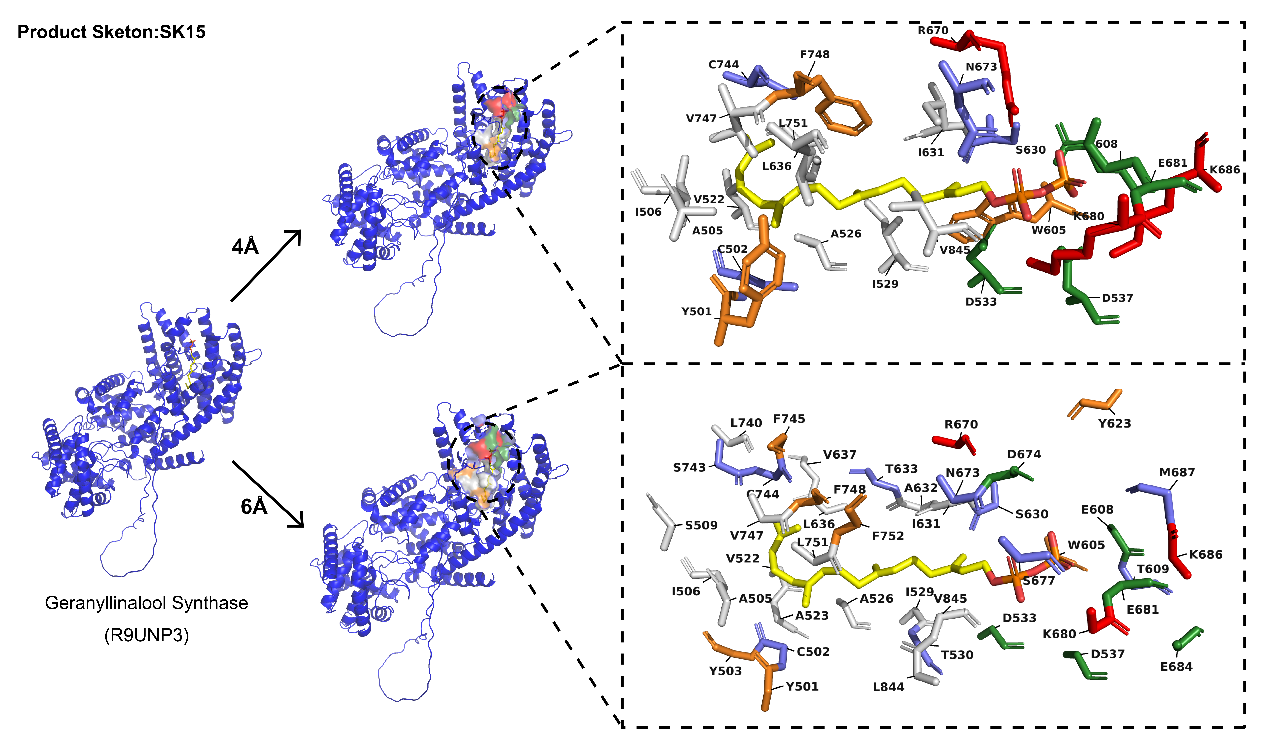

**Fig. S17**. Specific residue structures and topology formed by residues at 4Å and 6Å radial distances from the substrate of PdiTPSs producing the SK15 skeleton type.

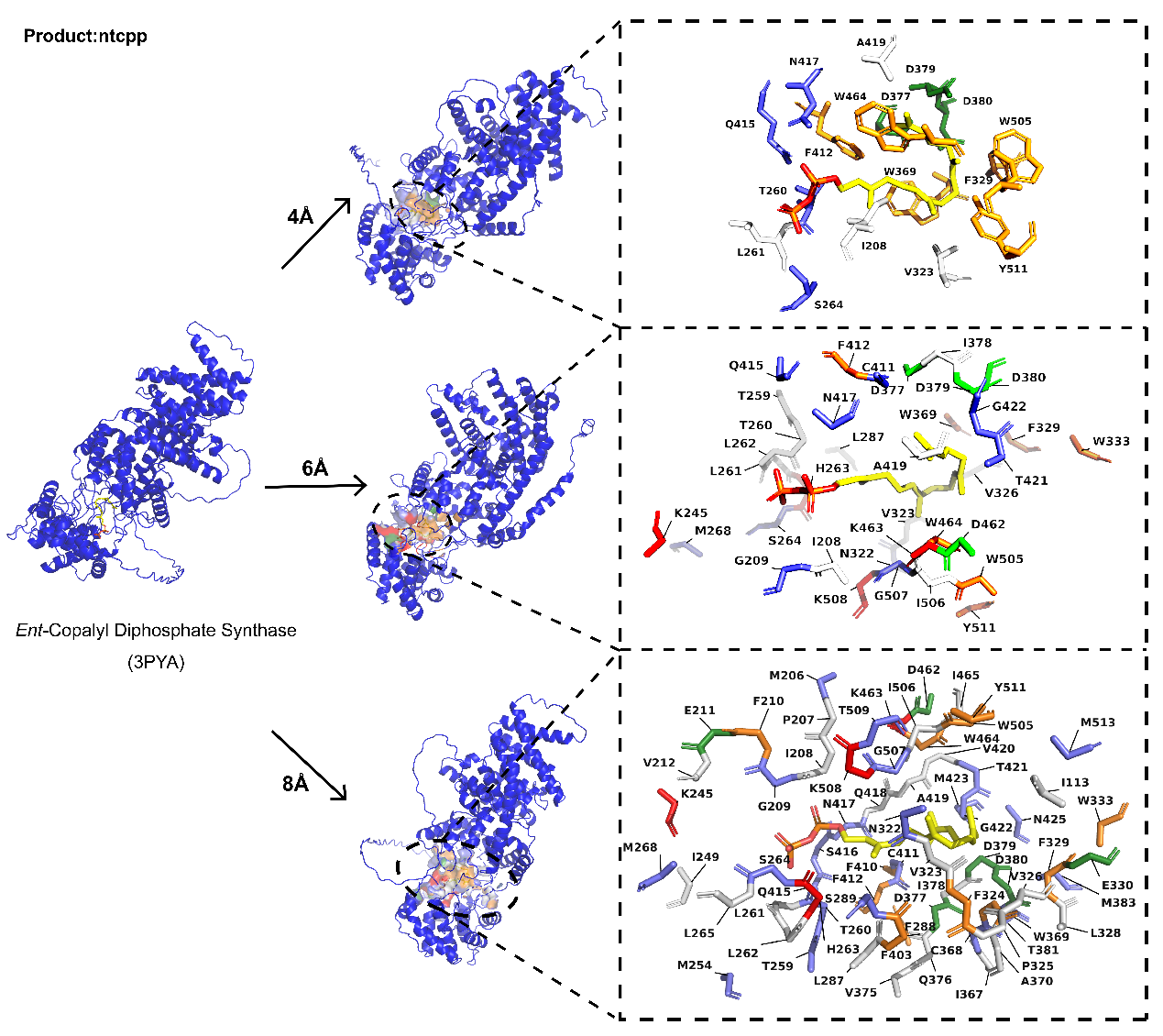

**Fig. S18**. Topological structure and specific residues within 4Å, 6Å, and 8Å radial of the substrate of PdiTPSs, which produce ntcpp and have confirmed functional residues.

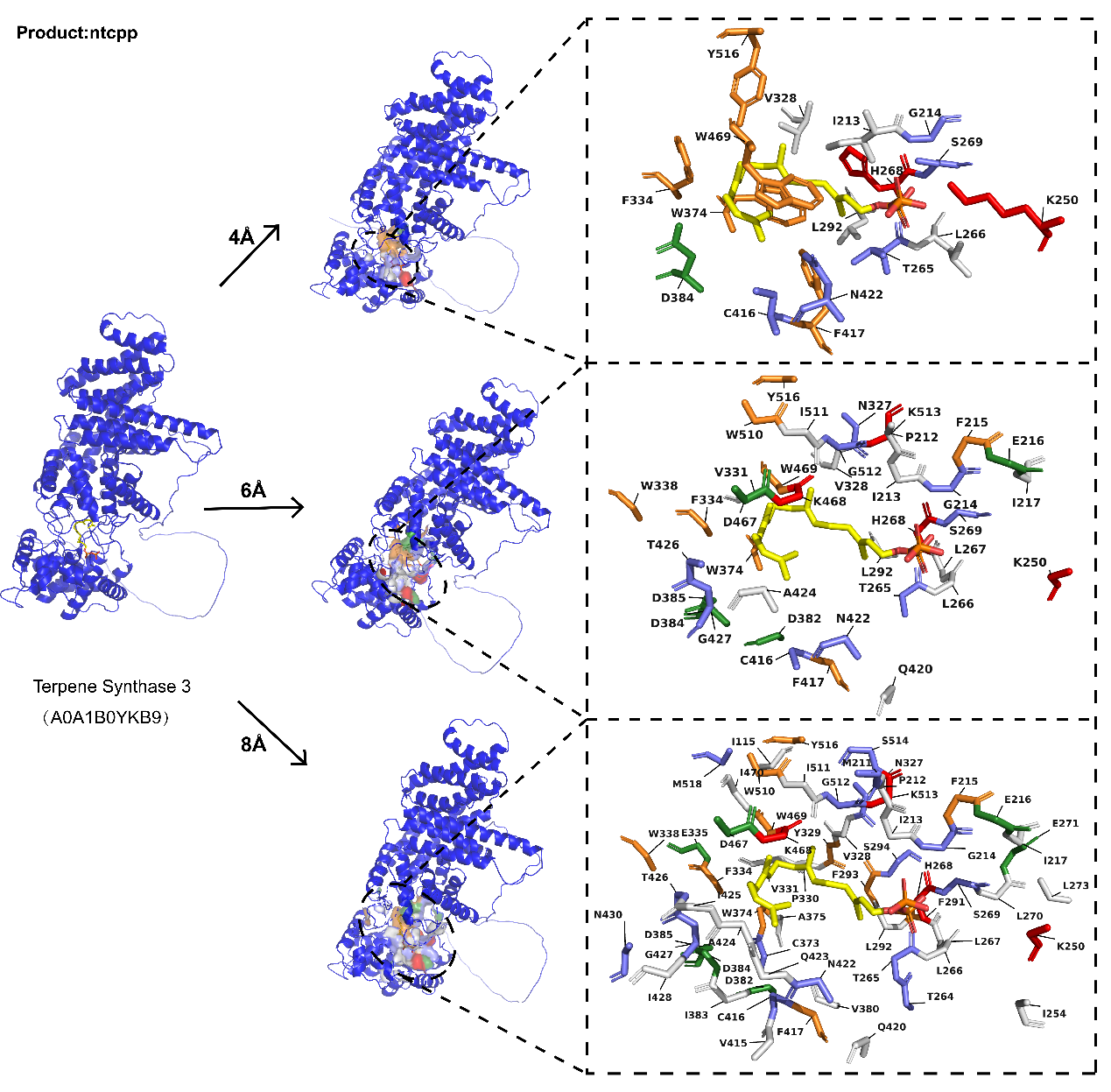

**Fig. S19**. Topological structure and specific residues within 4Å, 6Å, and 8Å radial of the substrate of PdiTPSs, which produce ntcpp and have confirmed functional residues.
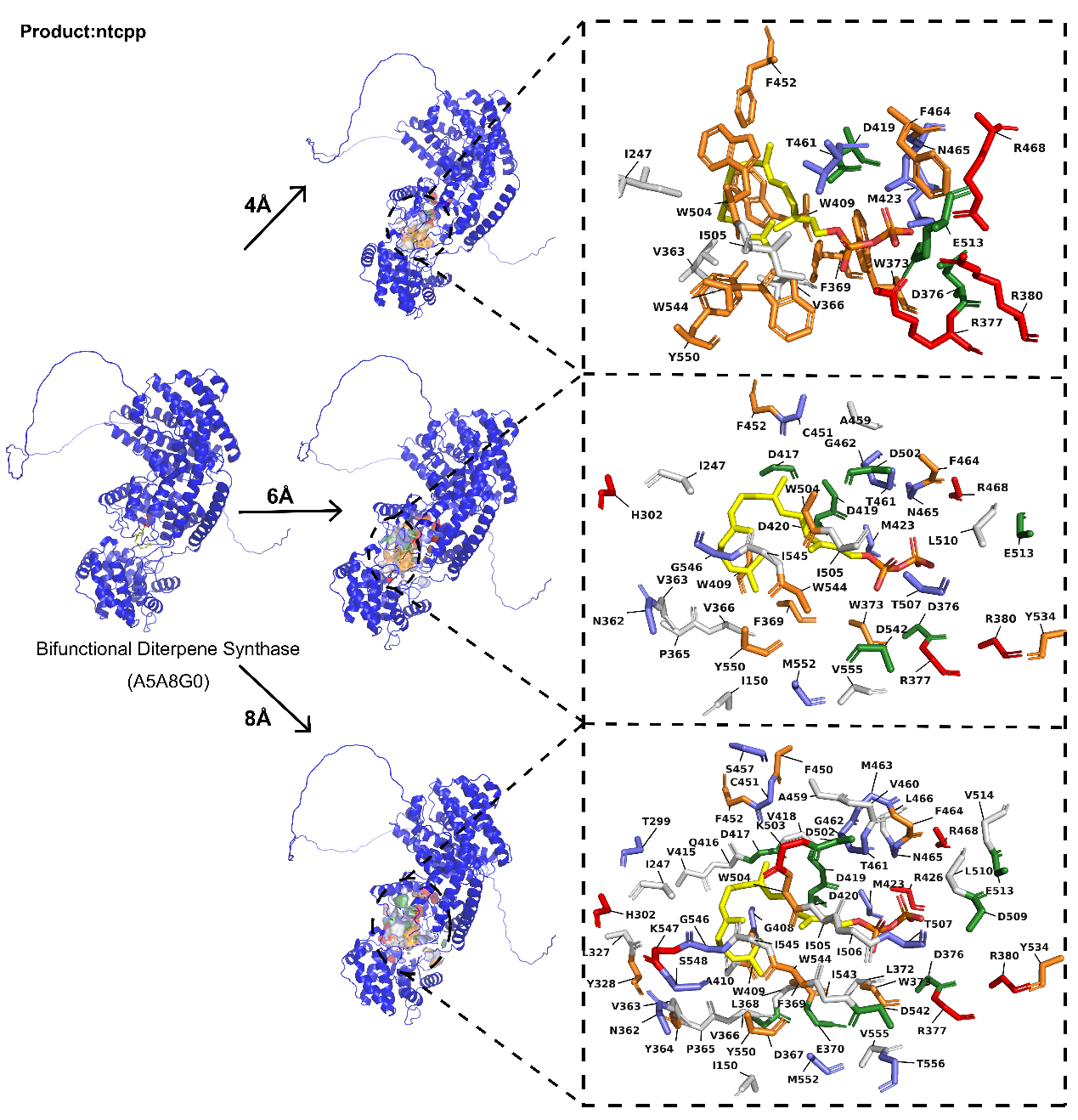

**Fig. S20**. Topological structure and specific residues within 4Å, 6Å, and 8Å radial of the substrate of PdiTPSs, which produce ntcpp and have confirmed functional residues.

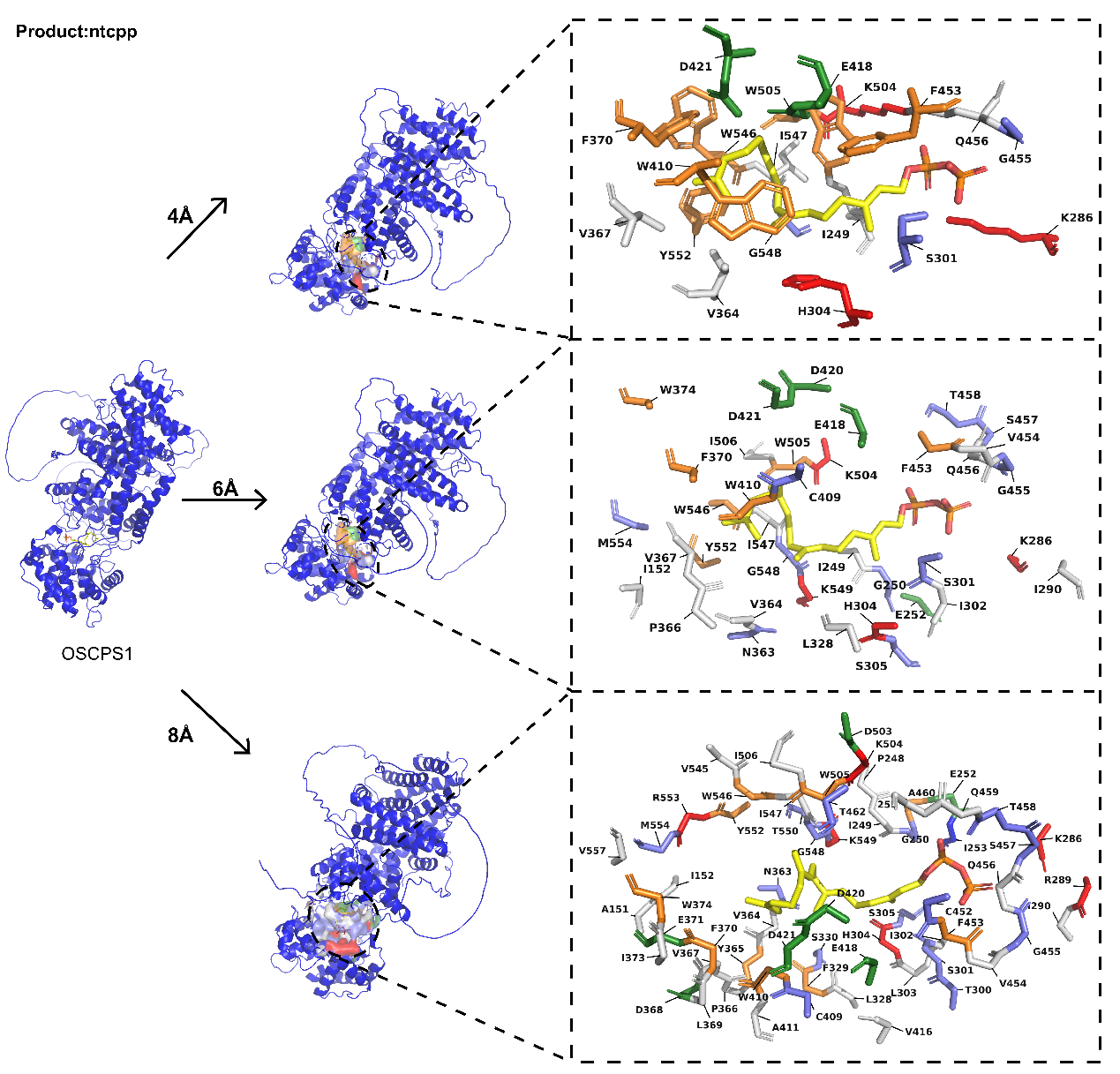

**Fig. S21**. Topological structure and specific residues within 4Å, 6Å, and 8Å radial of the substrate of PdiTPSs, which produce ntcpp and have confirmed functional residues.
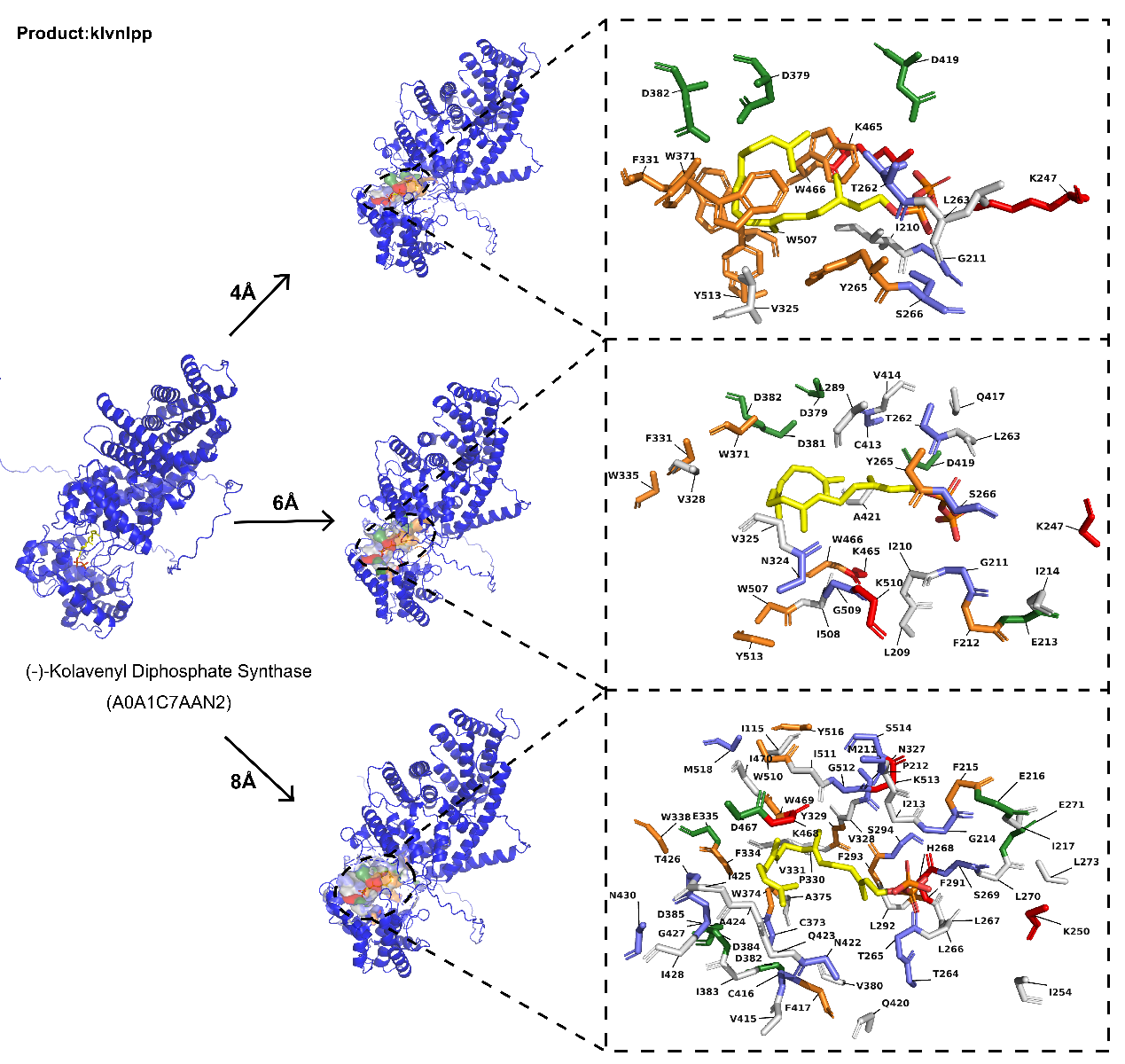

**Fig. S22**. Topological structure and specific residues within 4Å, 6Å, and 8Å radial of the substrate of PdiTPSs, which produce klvnlpp and have confirmed functional residues.
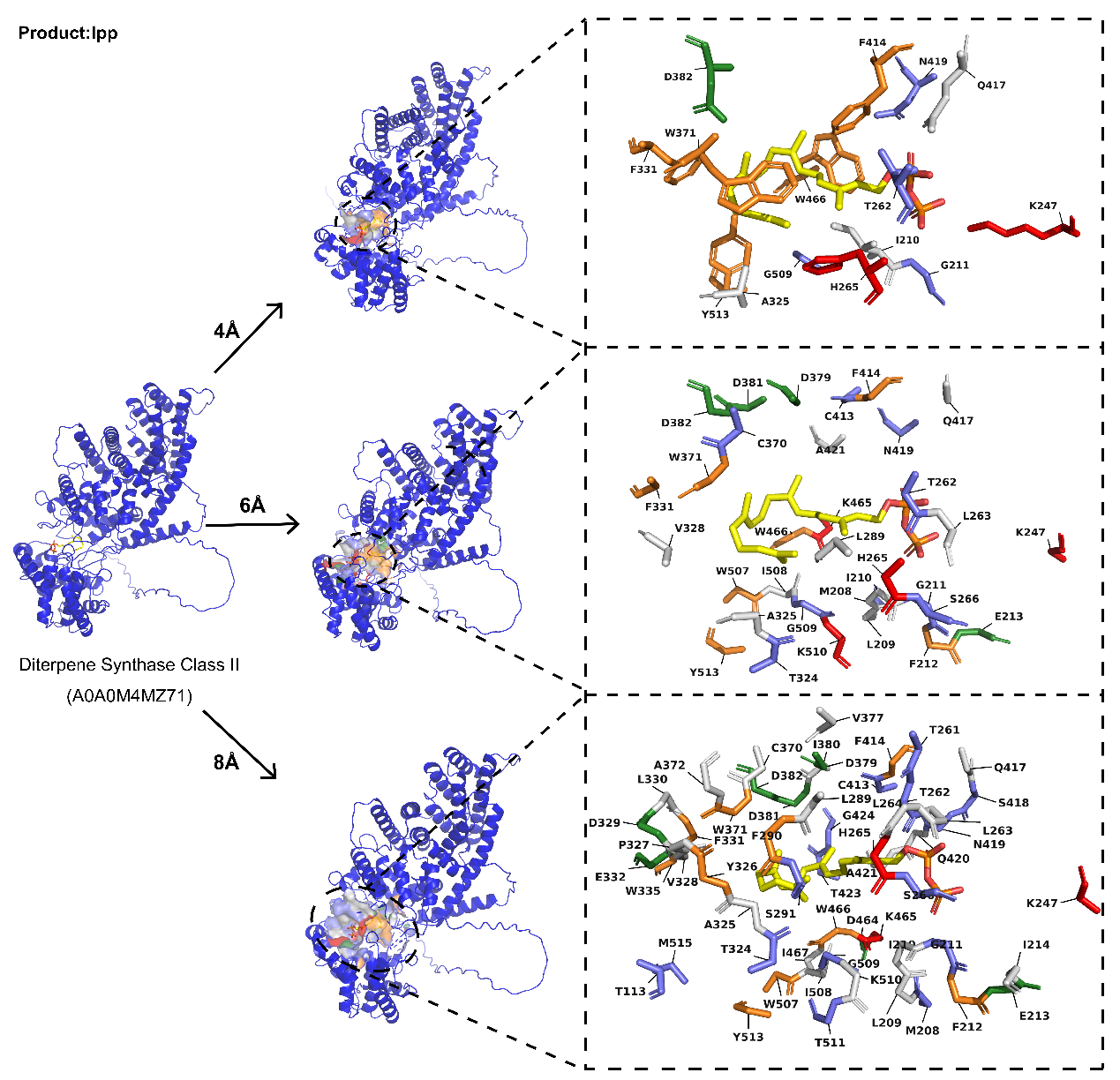

**Fig. S23**. Topological structure and specific residues within 4Å, 6Å, and 8Å radial of the substrate of PdiTPSs, which produce lpp and have confirmed functional residues.

**Supplementary Table**

**Table 5** Pearson's correlation coefficients between different sequence types and structures of PdiTPSs

| Content | Pearson's correlation coefficients | |
| --- | --- | --- |
| Nsequence : 4 Å radial | | 0.4 |
| Nsequence : 6 Å radial | | 0.36 |
| Nsequence : 8 Å radial | | 0.34 |
| Nsequence : 10 Å radial | | 0.33 |
| Nsequence : Overall structure | | 0.63 |
| Csequence : 4 Å radial | | 0.55 |
| Csequence : 6 Å radial | | 0.55 |
| Csequence : 8 Å radial | | 0.52 |
| Csequence : 10 Å radial | | 0.54 |
| Csequence : Overall structure | | 0.34 |
| NCsequence : 4 Å radial | | 0.6 |
| NCsequence : 6 Å radial | | 0.59 |
| NCsequence : 8 Å radial | | 0.55 |
| NCsequence : 10 Å radial | | 0.56 |
| NCsequence : Overall structure | | 0.55 |
| Overall sequence : 4 Å radial | | 0.51 |
| Overall sequence : 6 Å radial | | 0.46 |
| Overall sequence : 8 Å radial | | 0.42 |
| Overall sequence : 10 Å radial | | 0.42 |
| Overall sequence : Overall structure | | 0.7 |

Note: All *p* values in the table's statistical results are < 0.001.
